# Dissemination and progression of pulmonary *Mycobacterium avium* infection in mouse model is associated with type 2 macrophage activation

**DOI:** 10.1101/2021.06.23.449642

**Authors:** Raymond Rosenbloom, Igor Gavrish, Kerstin Seidel, Igor Kramnik, Nicholas A Crossland

## Abstract

Pulmonary infections caused by the group of nontuberculosis mycobacteria (NTM), *Mycobacterium avium* complex (MAC), are increasing worldwide and a growing public health concern. Pulmonary granulomas are the hallmark of MAC lung infection, yet reliable correlates of granuloma progression and susceptibility in immunocompetent hosts are poorly defined. The development of mouse models that recapitulate the diversity of granulomas seen in MAC pulmonary disease in humans is crucial to study mechanisms of susceptibility in humans and for preclinical evaluation of therapeutics. Unlike widely used inbred mouse strains, mice that carry the mutant allele at the genetic locus *sst1* develop human-like pulmonary tuberculosis featuring well-organized caseating granulomas. These mice became instrumental in pre-clinical testing of novel interventions. In this study we tested whether the B6.Sst1S that carries the *sst1* mutant allele on standard B6 background develop more advanced pulmonary infection with NTM *M. avium spp. hominissuis (M.av)*. To assess pulmonary disease progression, we utilized traditional semi-quantitative histomorphological evaluation and fluorescent multiplex immunohistochemistry (fmIHC) in combination with whole slide imaging and digital image analysis. After infection with the laboratory *M.av* strain 101, the B6.Sst1S pulmonary lesions progressed 12 – 20 weeks post infection, although we did not observe the formation of necrotic granulomas during this interval. Using fmIHC, we determined that the disease progression was associated with a steadily increasing proportion of mycobacteria infected Arg1+ and double positive iNOS+/Arg1+ macrophages. The B6.Sst1S granulomas had a greater proportion of Arg1+ and double positive iNOS+/Arg1+ macrophages, and decreased T cell density, as compared to wild type B6 mice. Thus, the genetic composition of the B6.Sst1S mice renders them more susceptible to pulmonary *M.av* infection. In combination with more virulent clinical isolates of *M.av* these mice could provide an improved mouse model that recapitulates more severe pulmonary disease in humans. The Arg1 macrophage expression in this model combined with automated fmIHC could serve as a sensitive biomarker for the unbiased assessment of medical countermeasures against NTM infection.

## INTRODUCTION

The nontuberculosis mycobacteria (NTM) *Mycobacterium avium* complex (MAC) is an emerging infectious disease in humans that causes pulmonary disease with increasing prevalence worldwide <u>(1)</u>. MAC consists of at least 9 environmental organisms, with *Mycobacterium avium* (*M.av*) being among the most common respiratory isolates (2). While the increasing burden of MAC pulmonary disease is mostly documented in high-income countries, MAC prevalence may be underappreciated and misdiagnosed with Tuberculosis (TB) in settings with limited diagnostic capabilities (3). Misdiagnosis of these two infections may contribute to poor clinical outcomes since each requires unique drug regimens (3). MAC pulmonary disease carries a poor prognosis (five-year all-cause mortality greater than 25%), and the only medical countermeasure available is an 18-24 month multidrug regimen with high toxicity (3, 4). Research into the pathogenesis and immune response to MAC infection is necessary to both guide the development of novel medical countermeasures against this pathogen and related mycobacterial species.

MAC infection in humans usually causes either disseminated or pulmonary disease depending on the host’s immune competence. CD4+ T cells and the IL-12, Interferon gamma (IFNγ) axis are essential features of immunity, as disseminated MAC infection classically occurs in the setting of the Acquired Immune Deficiency Syndrome and Mendelian Susceptibility to Mycobacterial Disease <u>(5)</u>. Pulmonary MAC disease typically presents with apical fibrocavitary or nodular bronchiectatic disease in individuals over the age of 60. Mechanisms of susceptibility to pulmonary MAC infection, however, are multifactorial and incompletely understood <u>(5, 6)</u>. They are strongly associated with structural lung defects caused by chronic obstructive pulmonary disease and cystic fibrosis, systemic autoimmune rheumatic diseases, immunosuppressive drugs such as TNF-alpha blockers, and malnutrition (5). The pathological hallmark of MAC pulmonary infection is granulomatous inflammation (5). MAC-positive granulomas with areas of caseous necrosis, epithelioid and multinucleated CD68+ giant cells were found in the lungs of immunocompetent patients with bronchiectatic and cavitary forms of the disease(7, 8). These findings support a role of necrotizing granulomas in bronchial ulceration, and the progression towards bronchiectatic and cavitary forms of pulmonary disease(9). The development of novel medical countermeasures will therefore partly depend on further investigation of local mechanisms of susceptibility in pulmonary granulomas.

Well studied in *Mycobacterium tuberculosis* (*M.tb*), granulomas are dynamic and heterogeneous immune responses involving both innate and adaptive immune cells (10). The granuloma structure and microenvironment consist of diverse macrophage phenotypes and CD4+ T cell populations that likely contribute to host resistance or immunopathology depending on the granuloma’s inflammatory state (10, 11). The signals that drive granuloma formation have largely been defined as mediated by the Th1 immune response consisting of IFNγ and classical macrophage activation (M1) via the expression of inducible nitric oxide synthase (iNOS) <u>(12)</u>. Macrophage’s expression of iNOS allows for the generation of nitric oxide (NO) to eliminate intracellular pathogens (13). In addition, alternatively activated macrophages (M2) that express Arginase-1 (Arg1) have also been described in TB and NTM granulomas(14, 15). Although typically found with the Th2 driven immune response to helminth infections, Arg1+ macrophages may have immunomodulatory or pro-fibrotic roles in mycobacterial granulomas(15, 16). Recently, type 2 macrophage response was shown to contribute to the necrotization of mycobacterial granulomas in zebrafish model(17). However, current understanding of mechanisms that underlie the necrotization and organization of mycobacterial granulomas remains limited(18).

Mouse models have played an instrumental role in elucidating essential features of the host response to *M.av* infection. As in humans, formation of granulomas is primarily a protective mechanism of innate immunity against *M.av* in immunocompetent mice (19). Also paralleling the immune response in humans, CD4+ Th1 cells producing IFNγ primarily mediate the adaptive immune response to pulmonary mycobacterial infection (19, 20). Deficiencies in IFNγ producing CD4+ T cells result in increased susceptibility to disseminated *M.av* infection in mice (21). Mouse models therefore recapitulate a defining feature of the immune response in humans in which defects in Th1 CMI result in progressive disease. Further research into pulmonary MAC in immunocompetent mice may therefore improve our understanding of susceptibility in human hosts.

The susceptibility to NTM in immunocompetent hosts is less well studied compared to immunodeficient hosts. Mouse models, however, were instrumental in elucidating the genetic determinants of *M.av* susceptibility in immunocompetent hosts. Previous work has demonstrated protection against aerosol infection with *M.av* (strain 724) by the resistant allele of the *Nramp1*(22). Recent work, however, using the C3HeB/FeJ strain, which carries the *Nramp1* resistant allele demonstrated that aerosol infection with *M.av* (strain 2285) resulted in chronic progressive lung infection and the formation of necrotic granulomas (23) The inbred C3HeB/FeJ mice are immunocompetent but highly susceptible to virulent M. *tb*. Unlike other standard inbred mouse strains, pulmonary tuberculosis (TB) progression in this mouse strain is characterized by the formation of well-organized human-like granulomas with necrotic centers(24–26). This unique necrotic granuloma phenotype is controlled by a single genetic locus *sst1* (<u>*s*</u>uper<u>*s*</u>usceptibility to <u>*t*</u>uberculosis-1), which has been characterized in our previous studies(27–31). We have shown that this locus controls macrophage responses to TNF and the *sst1* susceptibility allele reduces macrophage stress resilience and increases type I interferon production(32–35). It has been shown to control host resistance to several taxonomically unrelated intracellular bacterial pathogens(32, 36, 37). However, its role in the control of pulmonary NTM infection has not been studied.

The congenic strain B6.Sst1S generated in our laboratory carries the *sst1* susceptible allele on the B6 genetic background(30), which is *Nramp1*^*S*^. Because these mice carry two known susceptibility loci, we hypothesized that they may provide a model of pulmonary NTM infections that better resembles pulmonary NTM infections in immunocompetent, but susceptible humans. This mouse model of pulmonary NTM infections could therefore be used both to study mechanisms of immunocompetent host susceptibility to pulmonary NTM infections. In addition, mouse models of pulmonary NTM infections can be utilized for preclinical evaluation of therapeutics, such as recent work demonstrating the protective effects of a subunit vaccine candidate against *M.av* infection(38).

The goal of our study was to characterize *M.av* pulmonary lesion temporal progression in the B6.Sst1S mice and reveal correlates of susceptibility specifically in pulmonary granulomas. Using fluorescent multiplex immunohistochemistry (fmIHC) and quantitative image analysis (IA), we characterized the dynamics of macrophage populations. We found that progression of pulmonary lesions in more susceptible mice is associated with the increasing proportion of macrophages expressing iNOS, Arg1 and double positive iNOS+/Arg1+ macrophages. The susceptible mice also exhibited larger, more progressive pulmonary lesions with characteristic Interferon beta (IFNβ) expression in infected and uninfected iNOS+ macrophages and mononuclear cells adjacent to the lesions. The *sst1* locus mediated a slowly developing susceptibility to *M.av* infection by 16 weeks post infection (wpi). The primary histological feature in these lesions was granulomatous pneumonia lacking caseous necrosis. Utilization of fMIHC coupled with quantitative IA afforded us the ability to detect biologically significant changes that were not discernible with more traditional histopathological approaches. These insights offer enormous potential to elucidate granuloma correlates of susceptibility to mycobacterial disease.

## METHODS

### Mice and Mycobacterium avium infection

22 B6.Sst1S (16 females and 6 males) mice were enrolled in the study to assess mycobacterial CFU and pulmonary histopathology following 10^6 or 10^7 *M.av* (Strain 101) intranasal inoculation at 4-, 6-, 8-, 10-, 12-weeks post infection (wpi). In addition, B6 WT (N=12) and B6.Sst1S (N=10) female mice were enrolled in a comparison experiment and infected with 10^5 *M.av* (strain 101) by left mainstream intrabronchial inoculation. Mice were sacrificed at three time points: 12-, 16-, and 21-wpi. One B6 WT mouse and one B6.Sst1S mouse from 12 wpi were excluded from analysis because the lungs were atelectatic. Additionally, one B6.Sst1S 16wpi sample was excluded because the lungs were contaminated with eosinophilic crystalline pneumonia. Therefore for analysis the following specimens were used: 12wpi, N=2 B6.Sst1S and N=3 B6 WT; 16 wpi, N=3 B6.Sst1S and N=4 B6 WT; 21wpi, N=3 B6.Sst1S and N=4 B6 WT were used for H&E histopathology, but due to technical challenges, N=2 B6.Sst1S and no B6 WT mice were used for mHIC analysis. Additionally, two B6.Sst1S - YFP mice were infected with 10^6 *M.av* (strain 101) by left mainstream intrabronchial inoculation and sacrificed at 23 wpi. The B6.Sst1S - YFP mice were generated by crossing B6.Sst1S with a reporter mouse that has a Yellow Fluorescent Protein (YFP) reporter was inserted downstream of the IFNβ promoter where YFP serves as a surrogate for Interferon beta (IFNβ) expression (39). Mice were deeply anesthetized intraperitoneally with a mix of ketamine/xylazine. After euthanasia, the mice were perfused through the retro-orbital sinus with PBS/Heparin using AutoMate in-Vivo Manual Gravity Perfusion System (Braintree Scientific) to remove blood from the lungs through the retro-orbital sinus. The retro-orbital sinus was chosen to prevent lung collapse during the perfusion process. These steps were followed by perfusion with 4% paraformaldehyde and the lung instillation with 1% agarose.

### Tissue Inactivation, Processing, and Histopathologic Interpretation

Tissue samples were fixed for 24 hours in 10% neutral buffered formalin, processed in a Tissue-Tek VIP-6 automated vacuum infiltration processor (Sakura Finetek USA, Torrance, CA), followed by paraffin embedding with a HistoCore Arcadia paraffin embedding machine (Leica, Wetzlar, Germany) to generate formalin-fixed, paraffin-embedded (FFPE) blocks, which were sectioned to 5 μm, transferred to positively charged slides, deparaffinized in xylene, and dehydrated in graded ethanol. A subset of slides from each sample were stained with hematoxylin and eosin (H&E) for histopathology.

Using a Leica RM2255 Microtome, 2-to-3 step sections ~1 mm apart of whole lung sections were examined for 12 and 16 wpi specimens and targeted depth was used for 21 and 23 wpi specimens based on specific MRI coordinates. All animals except 23 wpi B6.Sst1S - YFP specimens were scored by a single board-certified pathologist (N.A.C.) using a semi-quantitative ordinal scoring system (see Table 1).

**Table 1.** Ordinal Grading Score: *Mycobacterium avium* intranasal and intrabronchiolar inoculation.

|  |  |
| --- | --- |
| 0 | No significant findings observed. |
| 1 | Multifocal lymphohistiocytic perivascular and peribronchiolar inflammation; minimal granulomatous pneumonia(<10% parenchyma). |
| 2 | Multifocal lymphohistiocytic perivascular and peribronchiolar inflammation; mild-to-moderate granulomatous pneumonia with rare intralesional single cell necrosis and neutrophils (>10-30% parenchyma). |
| 3 | Multifocal lymphohistiocytic perivascular and peribronchiolar inflammation; moderate-to-marked granulomatous pneumonia with mild intralesional single cell necrosis and neutrophilic infiltrate, sporadic multinucleated macrophages) and mild intralesional pulmonary fibrosis (>30-50% parenchyma) |

### Fluorescent Multiplex Immunohistochemistry (fmIHC)

Fluorescent mIHC (fmIHC) was performed using Opal fluorophores, which utilize tyramine signaling amplification (TSA) (Akoya Biosciences, Marlborough, MA). One manual 6-plex mIHC panel and one automated 7-plex mIHC panel were developed, optimized, and applied on lung sections from all animals in the study. Target antigens and optimization details for the manual 6-plex are outlined in Table 2. All sections were fixed in 10% neutral buffered formalin for 20 minutes to help retain cellular structure prior to antigen retrieval. Heat induced antigen retrieval (HEIR) was conducted in a BioCare Medical DeCloaking Chamber. Primary antibodies protocol parameters including concentration and sequence order were determined through optimization experiments. Next, secondary antibodies conjugated with horseradish peroxidase and TSA conjugated Opal fluorophores were applied for development of primary antibodies. Following development of each fluorophore, heat induced epitope retrieval (HIER) was reapplied to strip the previously developed primary-secondary antibody complexes, and the process was repeated in an iterative fashion. After development of the final fluorophore, slides were counterstained with DAPI and mounted with ProLong Gold Antifade Mountant (Thermo Fisher Scientific, Waltham, MA). Isotype controls were evaluated to confirm the absence of nonspecific tissue binding and lung sections from uninfected controls to confirm the specificity of the mycobacterium antibody.

**Table 2.** Optimized conditions for manual mIHC (5-plex)

| Sequential Order | AR Conditions | Rabbit derived primary antibody (Ab) | Manufacture, Catalog # | Primary Ab titration | Primary Ab incubation length | TSA-conjugated Fluorophore | Manufacture Catalog # | Opal Titration |
| --- | --- | --- | --- | --- | --- | --- | --- | --- |
| 1 | 90 degrees for 15min | Arginase-1 (Arg1) | Cell Signaling D4E3M | 1:100 | 1 hour at room temperature | 570 | Akoya Biosciences, FP1488001KT | 1:100 |
| 2 | 90 degrees C for 15min | Inducible nitric oxide synthase (iNOS) | abcam ab15323 | 1:200 | Overnight at 4 degrees C | 480 | Akoya Biosciences, FP1500001KT | 1:200 |
| 3 | 95 degrees C for 40 minutes & AR9 buffer | CD68 | Fisher Scientific PA5-32330 | 1:100 | Overnight at 4 degrees C | 620 | Akoya Biosciences, FP1495001KT | 1:100 |
| 4 | 95 degrees C for 40 minutes | CD3e | ThermoFisher MA5-14524 | 1:100 | 3 hours at room temperature | 520 | Akoya Biosciences, FP1487001KT | 1:200 |
| 5 | 90 degrees for 15min | *Mycobacterium tuberculosis* | Biocare Medical CP140A | 1:200 | 1 hour at room temperature | 780 | Akoya Biosciences, FP1501001KT | 1:12.5 |

Additionally, a 7-plex, automated mIHC assays for 21 wpi specimens and 23 wpi B6.Sst1S - YFP specimens were performed using a Ventana DISCOVERY Ultra Autostainer (Ventana Medical Systems, Roche Diagnostics, Basel, Switzerland), using DISCOVERY reagents according to the manufacturer’s instructions. Target antigens and protocol details for the automated multiplexes are outlined in Tables 3 & 4. The Ventana Discovery Ultra platform was also utilized to generate monoplex DAB assays utilizing serial sections derived from 23 wpi B6.Sst1S – YFP lungs to target *M.av* and green fluorescent protein (GFP).

**Table 3.** Optimized conditions for automated mIHC (6-plex)

| Sequential Order | Pre-Treatment-duration | Rabbit derived primary antibody (Ab) | Manufacture, Catalog # | Primary Ab titration | Primary Ab incubation length and temperature | TSA-conjugated Fluorophore | Manufacture Catalog # | Opal Titration |
| --- | --- | --- | --- | --- | --- | --- | --- | --- |
| 1 | CC1-95- 64 minutes | Arginase-1 (Arg1) | Cell Signaling Technology D4E3M | 1:100 | 24 min 37° C | 570 | Akoya Biosciences, FP1488001K T | 1:100 |
| 2 | CC1-95- 64 minutes | Inducible nitric oxide synthase (iNOS) | Abcam ab15323 | 1:200 | 32 min 37° C | 480 | Akoya Biosciences, FP1500001K T | 1:100 |
| 3 | CC1-95- 32 minutes | CD68 | Abcam ab12512 | 1:100 | 60 min 37° C | 690 | Akoya Biosciences, FP1497001K T | 1:100 |
| 4 | CC2-95- 64 minutes | CD4 | Cell Signaling Technology 25229 | 1:50 | 60 min 37° C | 620 | Akoya Biosciences, FP1495001K T | 1:100 |
| 5 | CC1-95- 64 minutes | CD3e | Thermo Fisher MA5-14524 | 1:100 | 60 min 37° C | 520 | Akoya Biosciences, FP1487001K T | 1:100 |
| 6 | CC1-95- 64 minutes | Myco-bacterium *tuberculosis* | Biocare Medical CP140A | 1:100 | 60 min 37° C | 780 | Akoya Biosciences, FP1501001K T | 1:25 |

**Table 4.** Optimized conditions for automated YFP mIHC (5 plex)

| Sequential Order | Pre-Treatment-duration | Rabbit derived primary antibody (Ab) | Manufacture, Catalog # | Primary Ab titration | Primary Ab incubation length | TSA-conjugated Fluorophore | Manufacture Catalog # | Opal Titration |
| --- | --- | --- | --- | --- | --- | --- | --- | --- |
| 1 | CC1-95- 64 minutes | Arginase-1 (Arg1) | Cell Signaling Technology D4E3M | 1:100 | 24 min 37° C | 520 | Akoya Biosciences, FP1487001 KT | 1:200 |
| 2 | CC1-95- 64 minutes | Mycobacterium *tuberculosis* | Biocare Medical CP140A | 1:100 | 60 min 37° C | 570 | Akoya Biosciences, FP1488001 KT | 1:125 |
| 3 | CC1-95- 64 minutes | GFP | Invitrogen A11122 | 1:100 | 24 min 37° C | 480 | Akoya Biosciences, FP1500001 KT | 1:250 |
| 4 | CC1-95- 64 minutes | Inducible nitric oxide synthase (iNOS) | Abcam ab15323 | 1:200 | 32 min 37° C | 690 | Akoya Biosciences, FP1497001 KT | 1:75 |
| 5 | CC1-95- 64 minutes | CD3e | Thermo Fisher MA5-14524 | 1:100 | 60 min 37° C | 620 | Akoya Biosciences, FP1495001 KT | 1:100 |

### Slide Scanning and Image Analysis

Whole slide images (WSI) of mIHC slides were digitized at 200x using the Vectra Polaris Automated Quantitative Pathology Imaging System (Akoya Biosciences). Exposure times were determined via autoexposure on a region of interest (ROI) on terminal B6.Sst1S slides with copious target protein. Remaining slides were batch loaded and digitized using the same exposure time parameters. An unstained lung section exposed to experimental HIER conditions was also imaged to create an autofluorescence signal that was removed from mIHC WSI using InForm software 2.4.9 (Akoya Biosciences). Inform Software was also used to computationally spectrally unmix digitalized mIHC slides using a pre-generated spectral library.

Digitized WSI were analyzed using the Halo IA software v3.2 (Indica Labs, Corrales, NM). Each slide was initially manually annotated to remove tissue artifacts (tissue folds, dust, fluorescent precipitates, etc.) and in two slides, areas of eosinophilic crystalline pneumonia in the left caudal lung lobe as evidenced by regional over-expression of the Arginase-1 and confirmation of the correlated histologic phenotype by a pathologist (N.A.C.). Next, each slide was manually annotated to denote the left (inoculated) and right (un-inoculated) lung lobes. Threshold values for visualization of each fluorophore were modified to minimize background signal and maximize specificity of target proteins. These threshold values were standardized for 12 and 16 wpi specimens, while unique thresholds were set for 21 and 23 wpi samples acknowledging that later timepoints were performed using an automated system that resulted in different signal intensities. IA was performed using the Tissue Classifier (TC) (a train-by-example machine learning algorithm utilizing a random forest hierarchical decision tree approach), Area Quantification (AQ), and High-Plex Fl (HP) modules. All module threshold values and parameters were modified and consistent for 12 and 16 wpi specimens, while unique parameters were created for 21 and 23 wpi specimens given vastly different fluorophore intensities. TC was trained to recognize granulomatous lesions containing predominantly iNOS+ histiocytes, *M.av*, and T cells. A second TC algorithm was designed to specifically recognize areas of T lymphocyte aggregates, characterized by abundant DAPI and CD3+ staining in the absence of mycobacterium antigen or macrophage markers. Additionally, a third TC algorithm was designed to recognize iNOS+ or Arg1+ histiocytic within lesions, while excluding CD3+ and other mononuclear cells.

TC outputs included the total percentage of area with immunoreactivity based on the total tissue area examined, as well as area (μm^2^). Lesion specific analysis was only conducted in the un-inoculated lung lobe at 16 wpi using the following criteria defined by the authors. TC granulomas in the un-inoculated lobe were separated into groups based on lesion area. For each unique specimen, lesions equal to and above the 90th percentile by area were annotated as “large secondary lesions,” and lesions between the 50th and 60th% percentile (inclusive) were annotated as “small secondary lesions.” Supplementary Figure 1 shows representative annotated lesions.

AQ outputs included the total percentage of tissue with immunoreactivity based on the total tissue area examined, as well as total percent positivity for two or more fluorophores (overlapping or non-overlapping fluorophores). AQ analysis was conducted on the whole lung lobe level and within the annotated uninoculated lung lobe lesions, as described above. HP outputs rely on cell segmentation via DAPI nuclear segmentation. The HP module was used to segment CD3+ cells (T cells) to determine density of T cells (# of CD3+ cells in the given area/ area analyzed μm^2^). The HP module and T cell density analysis was conducted on the whole lung lobe level, secondary lesions, and in secondary lesions that excluded TC classified lymphocytic aggregates.

### Statistical Analysis

For ordinal scoring, Kruskal-Wallis test with Benjamini, Krieger, and Yekutieli correction for multiple comparisons was used given the nonparametric, noncontinuous nature of the data. For comparison of two groups of ordinal histopathology scores, the Mann-Whitney test was used. For mIHC IA outputs, one-way analysis of variance with post hoc Tukey correction for multiple comparisons and unpaired student T-test were performed. All statistical analyses were performed using GraphPad Prism version 8.4.3 (GraphPad Software, La Jolla, CA) and for all tests, P < 0.05 was considered statistically significant.

## RESULTS

### *Mycobacterium avium* model of chronic infection in B6.Sst1S mice

To determine whether chronic disease can be induced after respiratory infection of B6.Sst1S mice with *Mycobacterium avium* strain 101 (*M.av*), we delivered 10^6^ or 10^7^ CFU of *M.av* intranasally. We observed progressive growth of mycobacteria in the lungs of the B6.Sst1S mice between 4- and 8-10-weeks post infection (wpi) and plateauing of the bacterial loads by 12 wpi. (Figure 1A). Among the mice that received 10^6^ CFU, there were statistically significant increases in the lung histology ordinal score at each timepoint (8, 10, 12 wpi) compared to 4 wpi (P=0.0127, P=0.0009, P=<0.0001 respectively) (Figure 1B).

**Figure 1.**
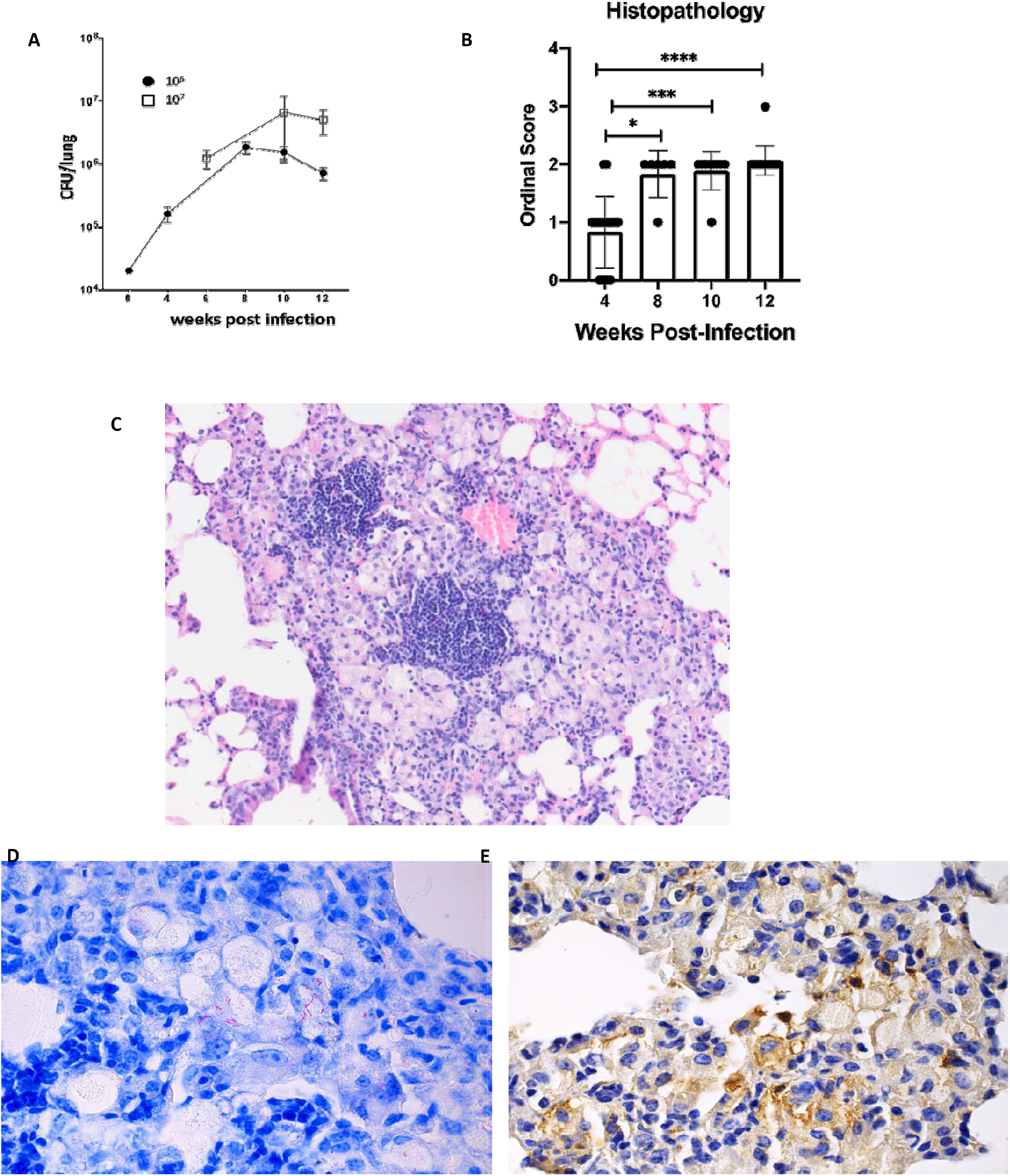
Establishment of chronic pulmonary M. *avium* infection with intranasal inoculation 4-12 weeks post infection (wpi) in B6.Sst1S mice. **A:** CFU/lung following intranasal (IN) 10^6 and 10^7 CFU IN infection. **B:** Histopathology scores of 10^6 CFU IN M. *avium* inoculated mice at 4, 8, 10, and 12 wpi. **C:** Representative micrograph of granulomatous lesion caused by M. *avium* at 12 wpi. **D:** Detection of M. *avium* Acid fast bacilli. **E:** Diaminobenzidine (DAB) immunohistochemistry detection of Mycobacterial antigen.For B, each data point represents a single tissue section. *P < 0.05, **P < 0.005, ***P < 0.0005, and ****P<0.00005. Original magnification, 100x (**C**) and 200x (**D** and **E**).

Microscopically, pulmonary lesions at 4 wpi were represented by minimal focal-to-multifocal lymphohistiocytic inflammation and perivascular lymphocytic cuffing with less than 10% of lung parenchyma affected. By 8 wpi, the lesions progressed to mild-to-moderate multifocal lymphohistiocytic inflammation that occupied 10-30% of the lung parenchyma. A board-certified pathologist (N.A.C.) utilized a semi-quantitative ordinal scoring system to assess lung pathology from each specimen (Supplementary Table 1).

At 12 wpi we observed moderate-to-marked multifocal lymphohistiocytic inflammation that affected >30-50% of lung parenchyma. At that stage we observed karyorrhectic debris (single cell necrosis), admixed with neutrophils and abundant swollen reactive histiocytes containing abundant lacy eosinophilic cytoplasm and perivascular lymphocytic cuffing (Figure 1C). Acid fast staining (Figure 1D) and IHC staining with antibodies targeting mycobacterial antigens (Figure 1E) demonstrated intracellular mycobacteria inside histiocytes within the lung lesions. However, neither extracellular bacteria, nor macro-necrosis or organized necrotic granulomas were observed.

These data demonstrate progression of pulmonary lesions and the establishment of chronic M.*av* infection in the lungs of the B6.Sst1S mice after respiratory challenge. In contrast to *M.tb* infection (30), however, no necrotic granulomas were observed in the lungs of the *M.av*-infected mice at 12 wpi.

### The development of primary bronchogenic and secondary lesions after unilateral intrabronchial inoculation

To enhance lung pathology, we utilized a model of intrabronchial inoculation with *M.av* where the bacterial inoculum is delivered to the left lung lobe. We analyzed tissue sections at 12, 16, and 21 wpi. The lesions in the inoculated lung lobes were characterized by extensive, multifocal granulomatous pneumonia, routinely emanating from peribronchial areas (Figure 2A). Despite extensive inflammation, the formation of necrotizing granulomas was not observed at the three time points examined (12, 16, 21 wpi).

**Figure 2.**
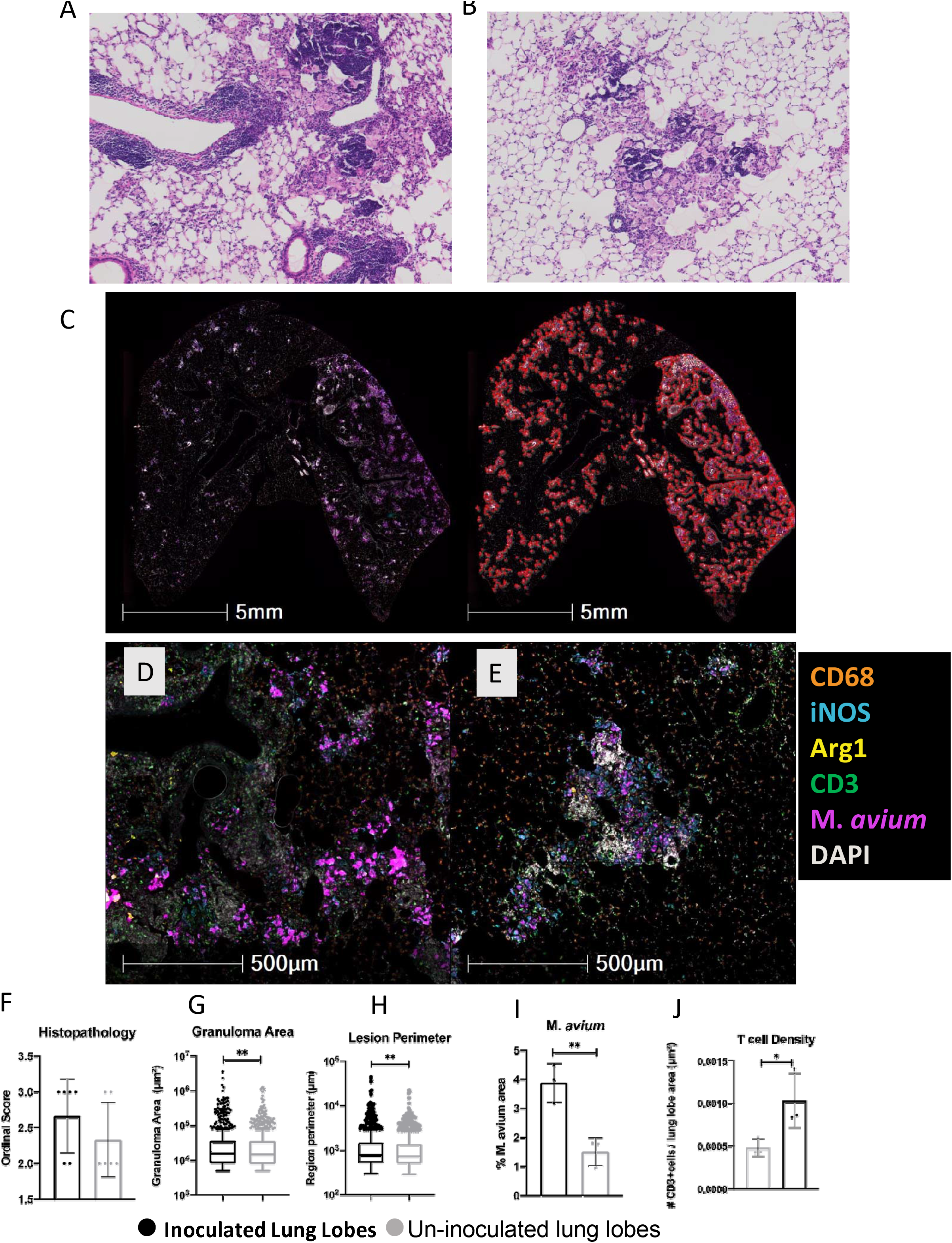
Comparison of primary and secondary lesions at 16 weeks-post infection (wpi) following unilateral left mainstream intrabronchial M. *avium* inoculation in B6.Sst1S mice. **A:** Representative micrograph of primary lesions in the inoculated left lung lobe with granulomatous infiltrate in immediate proximity to prominent peribronchiolar lymphoid infiltrate. **B:** Representative micrograph of secondary lesions in an un-inoculated right lung lobe composed predominantly of granulomatous infiltrate with lesser lymphoid aggregates. **C:** Representative whole lung raw fluorescent multiplex immunohistochemical (mIHC) (left) and random forest tissue classifier marked-up image (right) for detection of granulomatous lesions. **D:** Representative raw fluorescent mIHC image of primary lesions in the inoculated left lung lobe. **E:** Representative raw fluorescent mIHC image of secondary lesions in an un-inoculated right lung lobe. **F-J:** Inoculated vs un-inoculated lung lobes; **F.** Ordinal scores, **G.** lesion area, **H.** lesion perimeter, **I.** M. *avium* immunoreactivity area quantification (AQ), and **J.** T cell density. For F, each data point represents a single tissue section. For G-H, each data point represents a single lesion. For I-J, each data point represents a single animal. Data are expressed as means ± SD or with Tukey whiskers. Asterisks denote P values with Mann Whitney test (*) or Student’s unpaired T test (*_*). *P < 0.05, **P < 0.005. Original magnification, 25x (**C**), 50x (D), and 100x (**A** and **B**).

Unexpectedly, the uninoculated lung lobe also contained prominent granulomatous lesions at each time point examined. These lesions tended to have more random interstitial distribution with no direct affiliation with bronchioles and less evident perivascular inflammation (Figure 2B). At each time point examined (12, 16, 21wpi), ordinal histopathology scores tended to be increased in the inoculated compared to uninoculated lung lobes (Figure 2, Supplementary Figure 2). In summary, uninoculated lung lobe lesions could be clearly distinguished from the primary lesions in the left lobes: they were not associated with bronchi and were more organized with discrete foci of granulomatous pneumonia amongst larger areas of normal lung interstitium (Figure 2B). We concluded that these uninoculated lobe lesions, or “secondary lesions” resembled hematogenous spread, rather than the bronchogenic dissemination apparent in “primary lesions” in the inoculated lung lobes.

In order to further characterize the myeloid cell populations of these lesions, a fluorescent multiplex immunohistochemistry (fmIHC) was employed with antibodies specific for Mycobacterium, iNOS, Arg1, CD68, and CD3e. Macrophage phenotypes were distinguished using the markers CD68, iNOS, and Arg1, and T cells were identified with CD3e. Following fmIHC staining, whole slide images (WSI) were digitized for digital image analysis (IA), thereby facilitating unbiased quantitative analysis of the whole lung from each specimen.

To compare the primary lung lesions in the inoculated left lobe with the secondary lesions in the uninoculated lobe using fmIHC, we compared low magnification fmIHC images and detected salient differences between the lobes. Qualitative differences in degree of immunoreactivity for all biomarkers were markedly visible at 16 wpi (Figure 2C, *left*). We utilized a random forest Tissue Classifier (TC) for automated and unbiased identification of pulmonary lesions (see Methods). Classified primary lesions appeared more continuous and less discrete compared to secondary lesions (figure 2C, *right*). Quantitative IA demonstrated that primary lesions (inoculated lung lobes) tended to have larger lesion area (P value=0.0091) and perimeter (P value= 0.0031) than secondary lesions (un-inoculated lung lobes) at 16 wpi (figure 2G-H).

Further analysis of cell populations showed more differences between primary and secondary lesions. Primary lesions had qualitatively higher intensity *M.av* immunoreactivity within iNOS+ macrophages compared to secondary lesions (Figure 2D-E). There were also large areas of bronchial associated lymphatic tissue (BALT) with high density DAPI staining with CD3+ T cells within and around primary lesions (Figure 2D). Within secondary lesions, there was decreased area and intensity of *M.av* immunoreactivity (Figure 2E). Additionally, *M.av* immunoreactivity was more punctate and more easily visualized within iNOS+ macrophages (Figure 2E). The secondary lesions were not associated with bronchi and large areas of BALT were generally not observed in association with these lesions (Figure 2E). Quantitative analysis comparing inoculated and uninoculated lung lobes revealed that area of *M.av* immunoreactivity (P value= 0.0075) and T cell lesion density (P value=0.0464) was greater in the primary lesions within inoculated lobes compared to the secondary lesions in the uninoculated lung lobes at 16 wpi. (Figure 2 I-J) Increased histopathology score, mean lesion area, *M.av* and T cell immunoreactivity were consistently increased in the inoculated lung lobes at each time point examined (Supplementary Figure 2).

Taken together, histopathology and fmIHC demonstrated a common trend in which primary lesions could be distinguished from secondary lesions by enhanced pathology, association with bronchi, and increased mycobacterial immunoreactivity and greater T cell density.

### Progression of secondary lesions

After observing these disparate histopathological and immune signatures between primary and secondary lesions, further analysis was conducted to study secondary lesion temporal progression at 12, 16, and 21 wpi (Figure 3). At 12 wpi, secondary lesions were characterized by granulomatous pneumonia with sporadic single cell necrosis (Figure 3A). Alveolar macrophages became more reactive overtime represented by an increased area of lacy eosinophilic cytoplasm, with a greater overall total percentage of lung involvement (Figure 3B). Additionally, necrosis, frequency of multinucleated macrophages, and fibrosis became more frequent at later time points (Figure 3B). Although statistical significance was not reached, a trend toward increased ordinal histopathology scores was observed at later time points compared to 12 wpi (Figure 3G).

**Figure 3.**
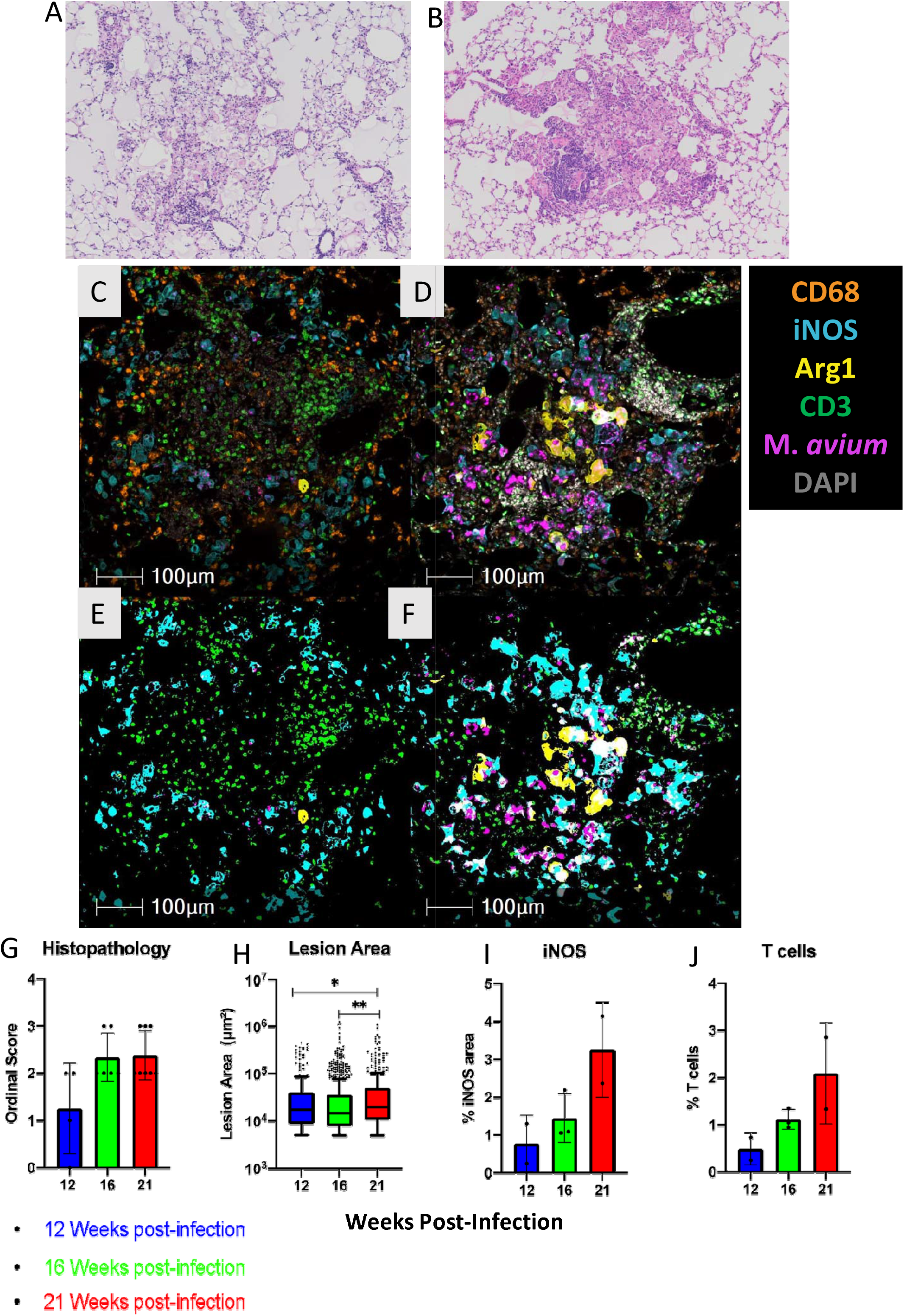
Progression of secondary lesions between 12-, 16-, and 21-weeks post infection (wpi) in un-inoculated right lung lobes following unilateral left mainstream intrabronchial M. *avium* inoculation in B6.Sst1S mice. **A:** Representative micrograph of early secondary lesion at 12 wpi. **B:** Representative micrograph of a large secondary lesion at 21 wpi containing abundant plump reactive macrophages. **C:** Representative raw fluorescent multiplex immunohistochemical (mIHC) image of small secondary lesion at 12 wpi. **D:** Representative raw fluorescent mIHC image of large secondary lesion at 16 wpi. **E:** Area quantification (AQ) work-up image of representative 12 wpi lesion (C). **F:** AQ work-up image of representative 16 wpi lesion (D). **G-J:** Un-inoculated lung lobes at 12, 16 and 21 wpi.; **G.** Ordinal scores, **H.** lesion area calculated using random forest tissue classification, **I.** iNOS immunoreactivity area quantification (AQ), **J.** CD3+ immunoreactivity AQ. For G, each data point represents one tissue section. For H, each data point represents one lesion. For I-J, each data point represents a single animal. Data are expressed as means ± SD or with Tukey whiskers. Asterisks denote P values with Kruskal-Wallis test (*) or one-way ANOVA (*-*). *P < 0.05, **P < 0.005. Original magnification 100x (**A-F**).

Analysis of cell populations within secondary lesions showed biomarker patterns associated with more advanced lesions. At 12 wpi, secondary lesions contained iNOS+ and iNOS-macrophages with few, sporadic Arg1+ macrophages (Figure 3C). There was punctate *M.av* immunoreactivity in some but not all iNOS+ macrophages (Figure 3C). T cells were also distributed in the lesion, generally congregated together in tertiary lymphoid aggregates (Figure 3C). At 16 wpi, a greater proportion of macrophages were iNOS+ and there was a greater presence of Arg1+ macrophages, including double positive iNOS+/Arg1+ macrophages (Figure 3D). In comparison to at 12 wpi, these iNOS+ and double positive macrophages appeared more reactive with greater cytoplasmic area (Figure 3D). Additionally, there was greater intensity and area of *M.av* immunoreactivity (Figure 3D). Quantitatively, there was a temporal progression of increased mean lesion area (12-21wpi P value=0.0488, 12-21wpi P value=0.0019), and percentage area of iNOS and T cells across the secondary lesion-containing uninoculated lung lobes at 12, 16, and 21 wpi. (Figure 3H-J).

Overall, secondary lesion temporal progression showed the development of larger lesions (Figure 3H). Given this trend, the smaller lesions likely represent newly formed lesions from hematogenous spread and de novo formation, whereas the larger lesions were more advanced, older lesions. We subsequently classified 16 wpi secondary lesions as “small” or “large,” as previously described in the Methods. Size stratification analysis allowed us to compare the temporospatial signatures of new and more advanced lesions, thereby facilitating the identification of macrophage correlates of granuloma progression.

Small secondary lesions tended to have less reactive macrophages with small cytoplasmic iNOS staining (Figure 4A). Arg1+ macrophages were generally not present and there were few T cells interspersed within the lesions (Figure 4A). In contrast, large secondary lesions contained more reactive iNOS+ and Arg1+ macrophages with larger cytoplasmic area (Figure 4B). Arg1+ macrophages were generally found in central parts of these large lesions (Figure 4B). There were also areas of high density DAPI staining with scattered CD3+ cells, which are likely tertiary lymphocytic aggregates (Figure 4B). Reactive macrophages as indicated by larger cytoplasmic areas found in large secondary lesions were uniformly CD68+ (Figure 4E) and largely iNOS+ (Figure 4F) with a smaller portion being Arg1+ (Figure 4G). These Arg1+ macrophages were frequently but not exclusively iNOS+ and mostly contained punctate *M.av* immunoreactivity (Figure 4H).

**Figure 4.**
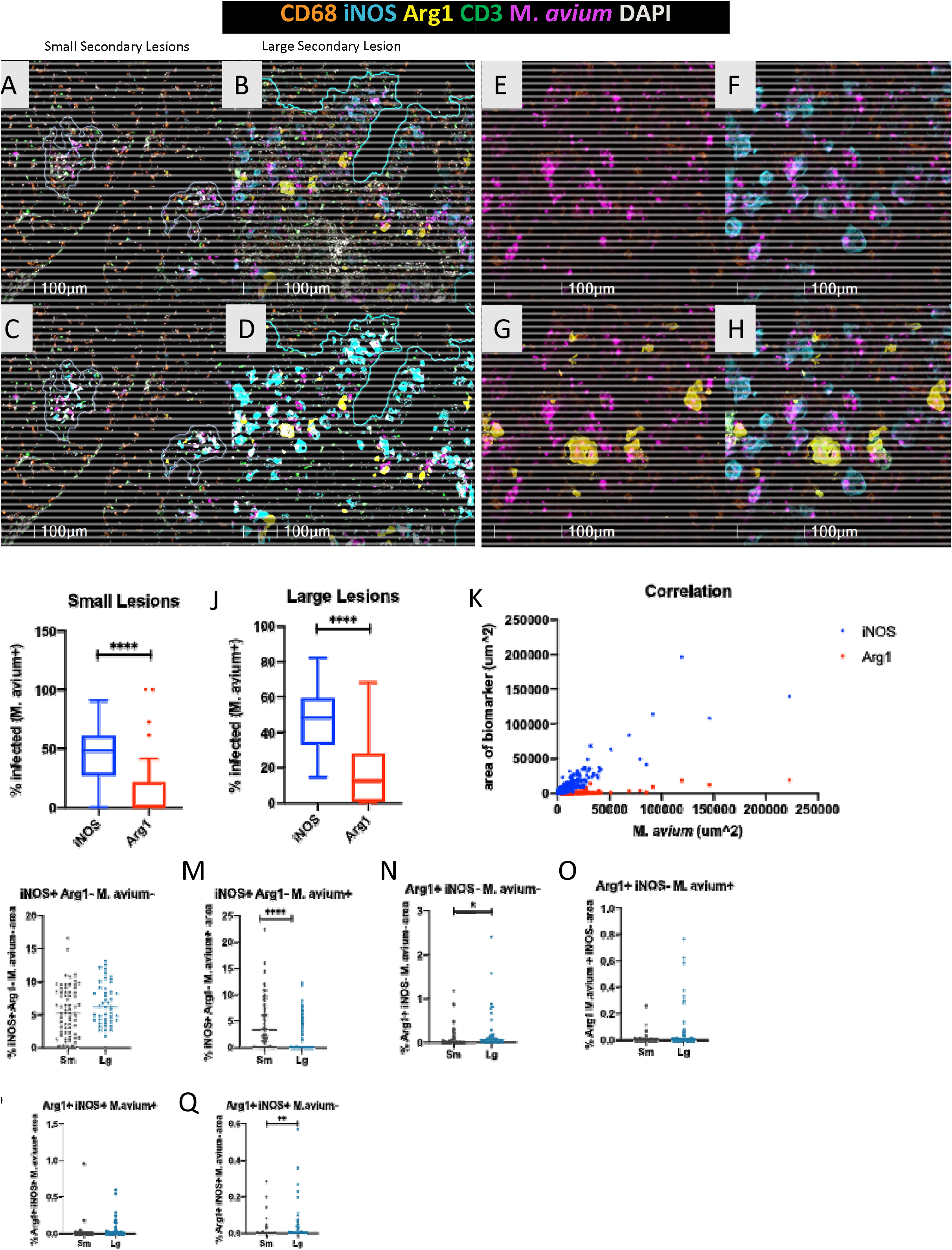
Characterization of macrophage populations in secondary lesions at 16 weeks-post infection (wpi) following unilateral left mainstream intrabronchial M. *avium* inoculation in B6.Sst1S mice. **A:** Representative raw fluorescent multiplex immunohistochemical (mIHC) image of small secondary lesion at 12 wpi. **B:** Representative raw fluorescent mIHC image of large secondary lesions at 16 wpi. **C:** Area quantification (AQ) marked-up image of representative small secondary lesion (A). **D:** AQ marked-up image of representative large secondary lesion (B). **E-F:** Macrophage phenotypes as visualized on raw fmIHC; **E.** CD68 and M. *avium* filers on, **F.** CD68, M. *avium*, iNOS filters on, **G.** CD68, M. *avium*, Arg1 filters on, **H.** CD68, M. *avium*, Arg1, iNOS filters on. **I:** Percentage antigen overlay of M. *avium*+ area with iNOS+ and Arg1+ in small lesions. **J:** Percentage antigen overlay of M. *avium*+ area with iNOS+ and Arg1+ in large lesions. **K:** Correlation between M. *avium* area on x-axis and iNOS or Arg1 area on y-axis. **L-Q:** Area quantification of iNOS, Arg1, and M. *avium* overlay in small and large granulomatous lesions; **L.** iNOS+/Arg1-/M. *avium*-area quantification (AQ). **M.** iNOS+/Arg1-/M. *avium*+ AQ, **N.** Arg1+/iNOS-/M. *avium*-AQ, **O.** Arg1/iNOS-/M. *avium*+ AQ, **P.** Arg1+/iNOS+/M. *avium*+ AQ, **Q.** Arg1+/iNOS+/M. *avium*-AQ. For I-Q, each data point represents a single lesion. Data are expressed on scatter dot plots with medians. Asterisks denote P values with Student’s unpaired T test. *P < 0.05, **P < 0.005, ***P < 0.0005, and ****P<0.00005. Original magnification 100x (**A-D**) and 200x (**E-H**).

Quantitative analysis of all secondary lesions revealed differential expression patterns of iNOS and Arg1 expressing macrophages. iNOS was more commonly associated with infection than Arg1+ macrophages. In small lesions, there was a greater percentage of iNOS macrophage immunoreactivity overlapped with *M.av* (45.75%) compared to Arg1 *M*.*av* overlap (14.43%) (P=<0.0001) (Figure 4I). This trend was the same in large lesions, with greater percentage iNOS M.*av* overlap (46.00%) compared to Arg1 *M.av* overlap (18.56%) (P=<0.0001) (Figure 4J). iNOS was strongly positively correlated with mycobacterial area (R squared= 0.8082; P=<0.0001), whereas the correlation between Arg1+ and *M.av* was lower (R squared= 0.5941; P=<0.0001) (Figure 4K). Percentage iNOS+ areas that did not overlap with *M.av* and Arg1 were similar in large and small lesions (Figure 4L). The proportion of iNOS+/*M.av*+ area was greater in small compared to large lesions (Figure 4M) (P=<0.0001). While iNOS remained the dominant macrophage phenotype, there was an increased presence of Arg1+ macrophages in large lesions. Large lesions contained a greater percentage of Arg1+ areas that were both *M*.*av* negative (P=0.0390) and *M.av* positive overlap compared to small lesions (Figure 4N-4O). These areas did not overlap with iNOS+ areas. Double positive iNOS+/Arg1+ macrophage immunoreactivity associated with *M.av* were predominantly identified in large lesions though statistical significance was not reached. (Figure 4P). When percentage area iNOS+/Arg1+ not overlapped with *M.av* was assessed, there was also increased percentage in large compared to small secondary lesions (P=0.0097) (Figure 4Q). The increased percentage of double positive iNOS+/Arg1+ areas not associated with infection also suggest that there was an increased proportion of uninfected double positive macrophages in large lesions. These quantitative findings demonstrating increased Arg1+ and double positive iNOS+/Arg1+ macrophages in large lesions are consistent with the qualitative observations previously described (Figure 4A-D).

Taken together, these data demonstrate that iNOS was the dominant infected cell phenotype in all small and large secondary lesions and thereby was strongly positively correlated with *M.av* infection. The appearance of infected and uninfected Arg1+ and double positive iNOS+/Arg1+ macrophages were specific findings associated with more advanced, larger lesions.

After characterizing these macrophage phenotype patterns associated with lesion progression, we were also interested in identifying myeloid and non-myeloid cells that expressed IFNβ. IFNβ has been identified as a possible correlate of susceptibility in human tuberculosis(40–42) and is implicated in *sst1-*mediated macrophage susceptibility (32–35). Therefore, determining the identity and spatial distribution of cells expressing IFNβ in mycobacterial lesions is crucial to delineate mechanisms of susceptibility. To answer this question, we used the B6.Sst1S, IFNβ-YFP reporter mouse in which YFP serves as a surrogate for IFNβ expression. We performed analysis at 23 wpi following *M.av* infection. To analyze representative lesions, we employed a fmIHC cocktail that included GFP specific antibodies, which served as the surrogate marker for IFNβ. Qualitative fmIHC demonstrated that IFNβ predominantly colocalized with iNOS+ *M.av+* macrophages. Additionally, IFNβ+ (iNOS-/Arg1-) cells with diffuse cytoplasmic expression were found within the perivascular mononuclear infiltrate and the pulmonary interstitium (Figure 5A). IFNβ+ cells were not typically found in T cell-rich lymphocytic aggregates (Figure 5B). There was no observable co-expression of Arg1+ macrophages and IFNβ in these lesions (Figure 5C). Overall, IFNβ expression was predominantly found in *M.av* infected macrophages, which were most commonly iNOS+. These findings were confirmed via serial sections using single marker staining with *mycobacteria-* and GFP-specific antibodies (Figure 5D-E).

**Figure 5.**
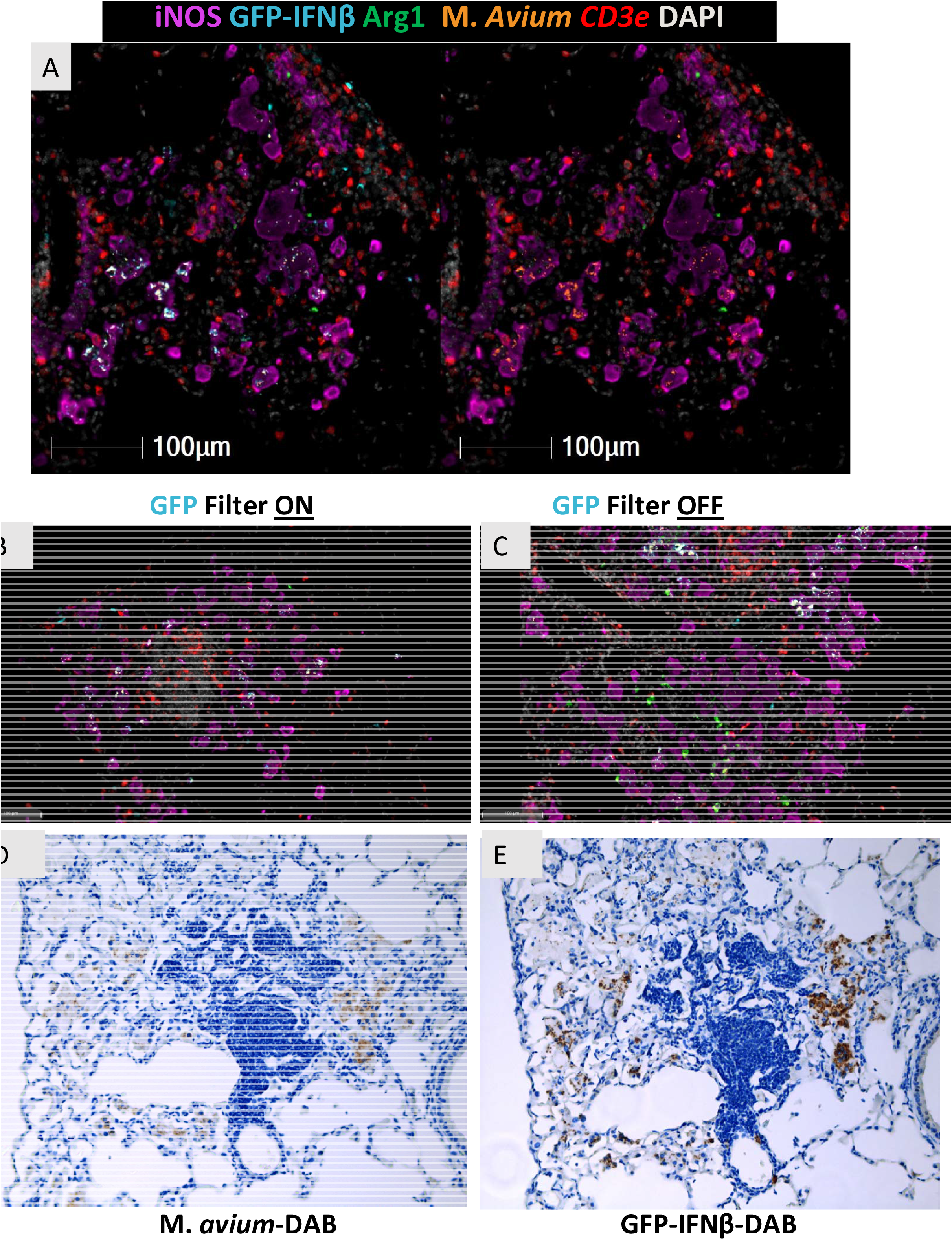
GFP-IFNβ expression in B6.Sst1S.YFP-IFNß mice intrabronchially inoculated with M. *avium* at 23 weeks post-infection (wpi). **A:** Representative raw fluorescent multiplex immunohistochemistry (mIHC) image of lesion with all filters on in the left panel and GFP filter off in the right panel. **B:** Representative raw fluorescent mIHC image of lesion with all filters on. **C:** Representative raw fluorescent mIHC image of lesion with all filters on. **D:** M. *avium*-Diaminobenzidine (DAB) IHC. **E:** GFP-IFNß DAB IHC stain. **D-E** are serial sections illustrating GFP-IFNß expression predominates in macrophages with M. *avium*. Original magnification 200X (**B-E**) or 400x (**A**).

### The *sst1*-mediated susceptibility to *M. avium* infection

To investigate the specific effect of the *sst1* locus in this model (B6.Sst1S) of *M.av* infection, we compared progression of pulmonary *M.av* with wild type B6 mice that carried the *sst1* resistant allele.

Histopathology and fmIHC image analyses were performed on B6 WT mice at 12, 16, and 21 wpi. Interestingly, ordinal histopathology scores obtained using H&E staining did not differ between WT and susceptible mice at individual timepoints (Supplemental Figure 3). Next, we applied quantitative unbiased whole lung lobe digital image analysis (IA) using both area quantification (AQ) and random forest tissue classifier (TC) as previously described. This analysis found no differences in lesion area of *M.av* immunoreactivity at 12 wpi (data not shown). At 16 wpi, there were no differences in lesion area or *M.av* immunoreactivity in the inoculated lung lobes (Figure 6A-B). However, there was a statistically significant decrease in percentage mean lesion area (P value=0.0373) and percentage *M.av* immunoreactivity (P value=0.0114) in the *sst1* resistant B6 WT uninoculated lung lobes (Figure 6C-D). This suggests that the Sst1 mechanism of resistance was exhibited in secondary lesions of the uninoculated lobes.

**Figure 6.**
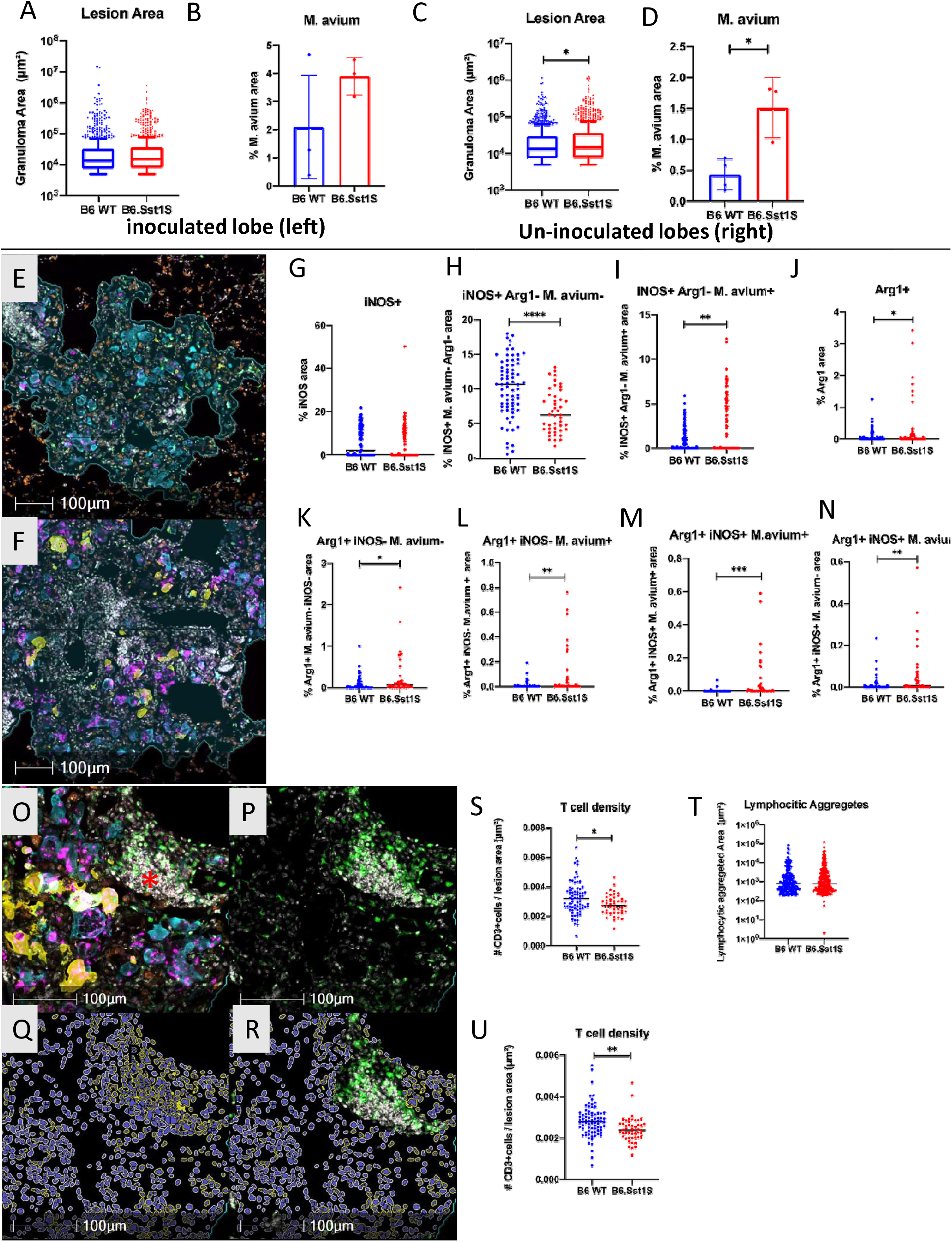
Comparison of immunopathology in infected and uninfected lung lobes from *M. avium* inoculated B6 WT compared to B6.Sst1S at 16 weeks-post infection (wpi) and the impacts of *sst1*-S in large secondary lesions at 16 wpi. **A-D:** inoculated (left) vs. un-inoculated lobes (right); **A.** lesion area, **B.** M. *avium* area quantification (AQ), **C.** lesion area, **D.** M. *avium* AQ. **E.** Representative raw fluorescent multiplex immunohistochemistry (mIHC) image of B6 WT large secondary lesion. **F:** Representative raw fluorescent mIHC image of B6.Sst1S large secondary lesion. **G-N:** iNOS, Arg1, and M. avium AQ immunoreactivity in B6 WT vs. B6Sst1S; **G.** iNOS AQ, **H**, iNOS+ Arg1-M. avium-AQ, **I.** iNOS+ Arg1-M. avium+ AQ, **J.** Arg1 AQ, **K.** Arg1+ iNOS-M. avium-AQ, **L.** Arg1+ iNOS-M. avium+ AQ, **M.** Arg1+ iNOS+ M. avium+ AQ, **N.** Arg1+ iNOS+ M. avium-AQ. **O:** Representative raw fluorescent mIHC image with all color filters on of secondary lesion with area of lymphocytic aggregation noted with (*). **P:** Raw fluorescent mIHC image (as in P) with only DAPI and C3+ filters on of secondary lesion with area of lymphocytic aggregation. **Q:** Highplex cell segmentation work-up image with yellow cytoplasmic cells representative CD3+ cells. **R:** Highplex cell segmentation work-up image (as in P) in areas excluding lymphocytic aggregates. **S**: T cell lesion density in B6 WT and B6.Sst1S large secondary lesions. **T:** Area of lymphocytic aggregates found in large secondary lesions of B6 WT and B6.Sst1S secondary lesions. Random forest tissue classifier was used to classify areas of lymphocytic aggregates (high density DAPI and CD3e+ staining) within large secondary lesions. **U:** T cell lesion density in B6 WT and B6.Sst1S large secondary lesions excluding lymphocytic aggregates. For B & D, each data point represents a single animal. For A, C, G-N, S-U, each data point represents a single lesion. Data are expressed on scatter dot plots with medians. Asterisks denote P values with Student’s unpaired T test. *P < 0.05, **P < 0.005, ***P < 0.0005, and ****P<0.00005. Original magnification 100X (**E-F**) or 200x (**O-Q**).

To investigate the differential cell populations in secondary lesions, we compared large and small lesions of B6.Sst1S and B6 WT mice. Both B6 WT and B6.Sst1S large secondary lesions were dominated by iNOS+ macrophages (Figure 6E-F). The proportion of iNOS macrophages was similar in the large lesions of resistant and susceptible mice (Figure 6G). Interestingly, however, small secondary B6 WT lesions contained marginally decreased percentage iNOS expression in comparison to B6.Sst1S (Supplementary Figure 3F) (P=0.034). The B6 WT lesions contained less *M.av* immunoreactivity compared to B6.Sst1S (Figure 6E-F). There was a greater proportion of uninfected iNOS+ macrophages in B6 WT large (P=<0.0001) and small (P=0.0009) secondary lesions, while there was a greater proportion of iNOS+ macrophages in B6.Sst1S large and small (P=<0.0001) lesions that were infected (Figure 6-H-I, Supplemental Figure 2G-H).

B6 WT large secondary lesions also contained lower percentage areas of Arg1+ (Figure 6J) (P=0.0176). There was a lower percentage of Arg1+ area either not associated with (P=0.0122) or associated with (P=0.0035) infection in large B6 WT lesions (Figure 6K–5L). A decreased proportion of double positive macrophages was also associated with the sst1R phenotype. Percentage areas of double positive iNOS+Arg1+ macrophages overlapped with *M.av* were decreased in large B6 WT lesions (P=0.0004) (Figure 6M). As previously discussed, the Arg1+ and double positive iNOS+Arg1+ macrophages were a unique feature of large well-developed lesions. We also compared the percentage area of iNOS+Arg1+*M.av*-negative areas to account for the decreased infection observed in B6 WT animals. This analysis also found decreased percentage iNOS+Arg1+ *M.av*-areas in large B6 WT lesions (P=0.0036), demonstrating that there were decreased proportions of double positive macrophages in these lesions compared to B6.Sst1S (Figure 6N) Therefore, both infected and uninfected double positive iNOS+Arg1+ macrophages were decreased in B6 WT large lesions. Interestingly, although as previously described histiocytic Arg1+ macrophages were characteristic of large lesions (Figure 4) and were qualitatively absent in most small lesions (Supplemental Figure 3D-E), quantitative analysis did detect decreased Arg1+ immunoreactivity in small B6 WT lesions compared B6.Sst1S lesions (Supplemental Figure 3J-M).

In addition to macrophage phenotypes, T cell density was also analyzed between large secondary B6 WT and B6.Sst1S lesions. T cell density was determined by first segmenting T cells using CD3e membranous staining and DAPI nuclear staining (Figure 5O-Q). T cell density was increased in B6 WT small (P=0.001) (Supplemental Figure 3N) and large (P=0.0131) secondary lesions (Figure 5S) compared to B6.Sst1S.

Large secondary lesions were frequently associated with areas of lymphocytic aggregates composed of iNOS-/Arg1-/CD68-likely representing tertiary lymphoid structures containing interspersed CD3e+ T lymphocytes (Figure 5O). These lymphoid structures were a similar area in B6 and Sst1S (Figure 5T). Using a tissue classifier, we removed these aggregates and analyzed T cell populations interspersed within mycobacterial lesions. Therefore, we conducted T cell density analysis following masking of these areas of lymphocytic aggregates (Figure 5R). Following this masking, T cell density remained greater in B6 WT lesions in comparison to B6.Sst1S (P=0.0025) (Figure 5U). These findings highlight that reduced T cell density was associated with granuloma susceptibility.

Taken together, the B6 WT sst1R phenotype was associated with reduced total *M.av* immunoreactivity and reduced average lesions area, as well as a lower proportion of Arg1+ and double positive iNOS+Arg1+ macrophages. Increased T cell density within lung lesions was also associated with the B6 WT sst1R phenotype. Combined, these patterns reflect slower progression of secondary mycobacterial lesions in the *sst1* resistant animals.

## DISCUSSION

To develop an improved mouse model and readouts for sensitive assessment of novel vaccines and therapeutics against pulmonary NTM infections, we pursued two interconnected goals: 1. To determine whether a mouse strain that carries two genetic loci that control susceptibility to intracellular pathogens including mycobacteria, *Nramp1* and *sst1*, represents a useful mouse model of pulmonary NTM infection in humans; and 2. To determine whether macrophage polarization markers may provide quantifiable correlates of the pulmonary NTM infection progression in this model.

Previous studies using experimental *M.av* infections in mice demonstrated that the genetic background of the host, route and dose of infection and M.*av* strains, laboratory or recent clinical isolates, all play an important role in establishing sustainable infection and determining the disease severity(23, 43, 44). We have chosen to use a virulent laboratory *M.av* strain MAC 101 (Chester, ATCC 700898), because this strain displayed a stable and virulent phenotype in other mouse studies: i) it multiplied in vivo and the infection was not completely resolved, ii) it induced T cell responses and inflammatory cytokine production, and iii) promoted inflammatory cell recruitment and pulmonary granuloma formation (21). This strain is also a standard strain for the evaluation of drug susceptibility in vitro and in vivo (43). In standard immunocompetent mouse strains B6 and BALB/c the respiratory infection with this strain produced stable infection that was self-contained within 6-12 weeks post infection (WPI) (43). Yet, in more than half of untreated human patients diagnosed with pulmonary MAC infection, the disease is progressive leading to more extensive lung damage and fibrocavitary pathology (45). This disease type is particularly unresponsive to antibiotic treatment. Therefore, a mouse model that recapitulates this form would be especially useful for testing combinations of antimicrobials and host-directed therapies targeting mechanisms underlying lung tissue damage.

We decided to use a mouse that carries the *sst1* susceptible allele, because in our previous studies this genetic locus was shown to specifically control the extent of lung pathology during infection with virulent M. *tb,* (30, 46), *Chlamydia pneumoniae*(32) and *Legionella pneumophila* (Ji, submitted for publication). Moreover, the C3HeB/FeJ that carries the *sst1*^*S*^ exposed to an aerosol infection with clinical isolate MAC 2285 produced progressive lung infection and pulmonary granulomas with areas of micronecrosis 6 weeks post infection (wpi) (23). However, the MAC infection in this study plateaued 7–9 wpi. The C3H mice carry the resistance allele of the *Slc11a1* (*Nramp1*) locus. This gene encodes a heavy metal pump in phagocytic vacuole and has been identified as a major genetic factor controlling replication of taxonomically unrelated intracellular pathogens, residing inside phagocytic vacuoles (47). It has been shown to control host macrophage resistance to *M.av* in vitro (48) and in a mouse model in vivo(22). The B6.Sst1 carries the susceptibility allele of the *Scl11a1*. In addition, it carries the MHC class II allele b of the H2-Aβ that has been also associated with increased lung pathology in *M.av*-infected mice (49). Therefore, we postulated that the genetic composition of the B6.Sst1S mice predisposes them to progressive pulmonary MAC infection and a more severe lung pathology, as compared to B6 and C3HeB/FeJ mice. We also extended the observation period to 12 – 20 wpi to assess progression and structural features associated with chronic pulmonary NTM infection.

The respiratory infection of the B6.Sst1S mice via intranasal inoculation with 10^7 CFU *M.av* inoculation resulted in chronic infection that was not resolved by the 12th week. At this time point, we documented the development of chronic interstitial lesions that contained macrophages loaded with mycobacteria (Figure 1). However, unlike *M.tb* infection, we found no organized necrotic granulomas in the lungs of the B6.Sst1S mice. To boost the infection, we employed a unilateral intrabronchial inoculation of *M.av.* In that model we observed the formation of chronic lesions in the inoculated lung lobe predominantly associated with bronchi. Interestingly, the contralateral uninoculated lung lobes also contained *M.av*-associated discrete lesions. At 12 wpi these lesions were smaller than the primary lesions in the inoculated lobe, and they were interstitial and not directly associated with bronchioles. Most likely, these lesions represent secondary, or metastatic, granulomas induced by the hematogenous spread of mycobacteria. Analysis of the discrete secondary lesions afforded us the ability to observe temporal patterns associated with the lesion progression. In addition to temporal analysis, we also stratified secondary lesions into small and large lesions at 16 wpi. We assumed that small lesions likely represented recently formed “early lesions,” whereas larger lesions were more advanced or “late lesions.” In addition to temporal progression, this lesion stratification afforded us the ability to identify correlates of granuloma progression within the same animal.

Temporal progression of the metastatic, secondary, lesions was associated with increased mean lesion area. At later time points, these lesions contained more *M.av*.-reactive, presumably infected, macrophages. Significantly higher proportion of the *M.av*.-infected macrophages expressed iNOS, than Arg1 at all timepoints. However, at the later time points, we observed an increased proportion of Arg1+ or double positive iNOS+/Arg1+ macrophages with *M.av*+ cytoplasmic staining. Similarly, at 16 wpi, larger secondary lesions contained an increased proportion of Arg1+ and double positive iNOS+/Arg1+ macrophages with cytoplasmic *M*.*av* immunoreactivity, as compared to smaller lesions. Thus, both comparisons based on either temporal or spatial criteria, demonstrate that more advanced lesions were associated with increased proportion of the Arg1+, and double positive iNOS+/Arg1+ macrophages.

We applied this metric to compare macrophage phenotypes between lesions in B6.Sst1S and B6 WT mice at 16 WPI. At this timepoint, the *sst1* effect on the lung lesion progression was not discernible using traditional semi-quantitative histomorphologic evaluation. However, using the fluorescent multiplex immunohistochemistry (fmIHC) criteria, the B6.Sst1S secondary lesions were classified as more advanced: they contained an increased fraction of *M.av*-infected macrophages, and an increased proportion of Arg1+ and double positive iNOS+/Arg1+ macrophages among the infected macrophages. Interestingly, the proportion of uninfected Arg1(+)/iNOS(-) macrophages was also increased in the B6.Sst1S granulomas, as compared to B6 WT. Taken together our studies using temporal, spatial, and genetic susceptibility criteria, demonstrated that increased Arg1 expression by macrophages in chronic *M.av* lesions represents a conserved macrophage biomarker of granuloma progression. Importantly, the appearance of double positive iNOS+/Arg1+ macrophages demonstrate that a binary M1/M2 macrophage polarity paradigm is insufficient to characterize the in vivo dynamics of macrophage activation within mycobacterial granulomas.

The Arg1+ macrophages were consistently detected in granulomas caused by various mycobacterial species and in various animal hosts. However, mechanisms of their induction and functional significance of Arg1+ macrophages in mycobacterial granulomas remain to be fully elucidated. The IHC analysis of human and non-human primate (NHP) TB lesions found macrophages expressing both iNOS and Arg1 enzymes(15, 50). In NHP granulomas, the Arg1+ macrophages were found primarily in the outer layer suggesting their role in limiting lung pathology (11, 15). A prominent role of the Arg1+ macrophages in the formation of fibrotic capsule was demonstrated using *Shistosoma mansoni*-induced granuloma(14). In an IL-13 overexpressing transgenic mouse, the infection with virulent M. *tb* resulted in the induction of Arg1+ macrophages and the formation of necrotic pulmonary granulomas driven by the arginase enzymatic activity (51). Similarly, the M2-like Arg1+ macrophage activation driving mycobacterial granuloma necrosis was observed in a zebrafish infected with *M.marinum*(17). The disease-promoting effects of Arg1+ macrophages in TB can be explained by arginine depletion and, thus, reducing NO production, a critical effector of host resistance to *M.tb* in mice (52). There is also experimental evidence that Arg1+ macrophages may also benefit the host. First, double iNOS and Arg1 knockout mice were found to be more susceptible to *M.tb* infection as compared to iNOS knockout mice, suggesting a protective role of Arg1 in these settings(53). Second, the previously cited Arg1+ roles in limiting inflammatory damage and formation of the fibrotic capsule suggest a role in granuloma organization. Unlike *M.tb* infection, iNOS deficiency is not associated with greater susceptibility to *M*.*av* infection in mice (20, 54). Thus, gradual accumulation of the Arg1+ macrophages during the chronic course of the *M.av* granuloma progression may represent an adaptive macrophage reaction limiting lung tissue damage by NO. Furthermore, depletion of L-arginine has been shown to suppress T cell proliferation and cytokine release (55) suggesting an Arg1 role in mitigating T cell-driven immunopathology (53). However, prolonged Arg1 expression may contribute to local immunosuppression and unresolving chronic infection. Of note, we observed reduced T cell density in advanced granulomas in our B6.Sst1S model. Although, a causal relationship between this phenomenon and the increase in Arg1 expressing macrophages remains to be established.

The signals that induce and maintain the Arg1 expression in macrophages of mycobacterial granulomas are not well characterized. Canonically, the M2 macrophage polarization and Arg1 expression are driven by Th2 lymphocyte-derived cytokines IL-4 and IL-13 (56). Mycobacterial infections, however, are known to drive predominantly Th1 responses (57, 58). Alternatively, mycobacteria were shown to induce Arg1 in macrophages directly via TLR – MyD88 – CEBPβ signaling. This pathway required no T cells, IL-4 or IL-13, nor STAT6 activation (16). Because the *sst1*-susceptible mice expressed higher levels of Arg1 in *M.av* lesions, we hypothesized that this locus may be involved in macrophage polarization as well. Our previous studies demonstrated that the aberrant macrophage activation underlies the *sst1* susceptible phenotype (32, 46). Upon activation, the mutant macrophages upregulate type I IFN (IFN-I) pathway that drives the integrated stress response (ISR) via upregulation of PKR and ATF4 and ATF3 transcription factors(33). Of note, the promoter for the Arg1 gene does include binding sites for ATF3. Therefore, we wanted to determine whether Arg1 in the B6.Sst1S background can be upregulated via the IFN-I-mediated ISR in a cell autonomous manner. We used B6.Sst1S-YFP-IFNβ knockin reporter mice that utilizes YFP as a surrogate for IFNβ and allows the identification of the IFNβ-producing cells in situ. By utilizing the fmIHC approach, we investigated co-localization of IFNβ/YFP with Arg1, iNOS and *M.av*. However, we did not observe significant co-localization of Arg1 and IFNβ within mycobacterial lesions. In contrast, IFNβ/YFP expression is predominantly colocalized with *M.av.-*infected iNOS+ Arg1-histiocytes. Further studies using this reporter mouse will allow characterization of the Arg1+ and IFNβ-producing cells, their interactions in situ and their respective roles in shaping granulomas induced by mycobacteria.

From the technological perspective, our work contributes to a growing body of infectious disease research showing the utility of whole slide imaging (WSI) platforms and digital image analysis (IA) tools to quantify morphomolecular and temporospatial signatures of multiple biomarkers in single tissue sections <u>(59, 60)</u>. In our findings, fmIHC and quantitative IA uncovered macrophage correlates of granuloma susceptibility and *Sst1S* susceptibility that were not observed with traditional histopathological approaches. Quantitative data outputs produced using WSI and digital IA facilitates more rigorous statistical analyses as compared to semi-quantitative ordinal methods used in traditional histopathology and mitigates limitations associated with interobserver variability.

Although the small number of animals analyzed at each timepoint is a limitation in this study, our digital IA approach enabled us to analyze hundreds of pulmonary lesions. Statistically significant results were observed both temporally, between mouse strains, and among large and small secondary lesions at 16 wpi. Another limitation is a narrow focus on macrophage M1/M2 markers. The integration of T cell markers in our spatial analysis will prove informative in linking local macrophage phenotypes with T cell responses within distinct granuloma compartments, as described by Gideon et al.(61). Technical limitations also include an impeded accuracy of segmentation of histiocytes due to their occasional multinucleation, as well as heterogeneity in cytoplasmic area and shape. This contrasts with the mononuclear CD3+ cells, which are relatively uniform in their overall size and shape and were easily segmented using DAPI nuclear staining. Therefore, to obtain quantitative outputs for the macrophage markers iNOS and Arg1, we utilized the Area Quantification (AQ) module to measure the area of these fluorescent stains. While this provides an accurate representation of the area of each biomarker and pixel overlay between multiple biomarkers, it does not directly phenotype individual cells. Despite these limitations, our approach provides uniformity of automated staining, whole slide scanning and unbiased digital image analysis. Therefore, it can be implemented on a larger scale in preclinical studies.

Because the structure of granulomas influences whether pharmaceutical compounds or a vaccine-induced immune response can penetrate granulomas to eradicate mycobacteria(23, 62, 63), the development of predictive mouse models that mimic the diversity of human pulmonary NTM disease is particularly important. Although we did not observe caseating necrotic granulomas in B6.Sst1S using laboratory adapted virulent *M.av* strain 101, previous work in C57BL/6 mice using clinical isolates has shown necrosis (19, 44). Our studies using advanced quantitative imaging and image analyses, demonstrated that B6.Sst1S mice are more susceptible to chronic *M.av* infection than B6 WT. Therefore, the utilization of B6.Sst1S and clinical isolates of *M.av* could provide an improved mouse model that recapitulates more severe forms of the disease in humans. Combined with quantitative imaging and Arg1 expression in macrophages as a biomarker associated with pulmonary NTM progression, this model will provide tools for cost effective and unbiased assessment of therapeutics and vaccines.

## ACKNOWLEDMENT

We would like to acknowledge expert advice of Drs. Keira Cohen (JHU) and Dr. Russ Huber (VA Jamaica Plain). This work was funded by R01 HL126066 and R01 HL133190 (I.K.). This work utilized a Ventana Discovery Ultra autostainer that was purchased with funding from NIH grant S10 OD026983 (PAR-18-600) (N.A.C.).

Authors declare no competing interests.

## Notes

### Competing Interest Statement

The authors have declared no competing interest.

## REFERENCES

1. Prevots DR, and Marras TK. Epidemiology of human pulmonary infection with nontuberculous mycobacteria: a review. Clin Chest Med. 2015;36(1):13–34.

2. Hoefsloot W, van Ingen J, Andrejak C, Angeby K, Bauriaud R, Bemer P, et al. The geographic diversity of nontuberculous mycobacteria isolated from pulmonary samples: an NTM-NET collaborative study. Eur Respir J. 2013;42(6):1604–13.

3. Baldwin SL, Larsen SE, Ordway D, Cassell G, and Coler RN. The complexities and challenges of preventing and treating nontuberculous mycobacterial diseases. PLoS Negl Trop Dis. 2019;13(2):e0007083.

4. Diel R, Lipman M, and Hoefsloot W. High mortality in patients with Mycobacterium avium complex lung disease: a systematic review. BMC Infect Dis. 2018;18(1):206.

5. Griffith DE, Aksamit T, Brown-Elliott BA, Catanzaro A, Daley C, Gordin F, et al. An official ATS/IDSA statement: diagnosis, treatment, and prevention of nontuberculous mycobacterial diseases. Am J Respir Crit Care Med. 2007;175(4):367–416.

6. Chin KL, Sarmiento ME, Alvarez-Cabrera N, Norazmi MN, and Acosta A. Pulmonary non-tuberculous mycobacterial infections: current state and future management. Eur J Clin Microbiol Infect Dis. 2020;39(5):799–826.

7. Fujita J, Ohtsuki Y, Shigeto E, Suemitsu I, Yamadori I, Bandoh S, et al. Pathological findings of bronchiectases caused by Mycobacterium avium intracellulare complex. Respiratory medicine. 2003;97(8):933–8.

8. Fujita J, Ohtsuki Y, Suemitsu I, Yamadori I, Shigeto E, Shiode M, et al. Immunohistochemical distribution of epithelioid cell, myofibroblast, and transforming growth factor-beta1 in the granuloma caused by Mycobacterium avium intracellulare complex pulmonary infection. Microbiology and immunology. 2002;46(2):67–74.

9. Kim TS, Koh W-J, Han J, Chung MJ, Lee J-H, Lee KS, et al. Hypothesis on the evolution of cavitary lesions in nontuberculous mycobacterial pulmonary infection: thin-section CT and histopathologic correlation. AJR American journal of roentgenology. 2005;184(4):1247–52.

10. Ramakrishnan L. Revisiting the role of the granuloma in tuberculosis. Nat Rev Immunol. 2012;12(5):352–66.

11. Flynn JL, Gideon HP, Mattila JT, and Lin PL. Immunology studies in non-human primate models of tuberculosis. Immunol Rev. 2015;264(1):60–73.

12. Pagán AJ, and Ramakrishnan L. The Formation and Function of Granulomas. Annual review of immunology. 2018;36(1):639–65.

13. Rath M, Muller I, Kropf P, Closs EI, and Munder M. Metabolism via Arginase or Nitric Oxide Synthase: Two Competing Arginine Pathways in Macrophages. Front Immunol. 2014;5:532.

14. Hesse M, Modolell M, La Flamme AC, Schito M, Fuentes JM, Cheever AW, et al. Differential regulation of nitric oxide synthase-2 and arginase-1 by type 1/type 2 cytokines in vivo: granulomatous pathology is shaped by the pattern of L-arginine metabolism. Journal of immunology. 2001;167(11):6533–44.

15. Mattila JT, Ojo OO, Kepka-Lenhart D, Marino S, Kim JH, Eum SY, et al. Microenvironments in tuberculous granulomas are delineated by distinct populations of macrophage subsets and expression of nitric oxide synthase and arginase isoforms. Journal of immunology. 2013;191(2):773–84.

16. El Kasmi KC, Qualls JE, Pesce JT, Smith AM, Thompson RW, Henao-Tamayo M, et al. Toll-like receptor-induced arginase 1 in macrophages thwarts effective immunity against intracellular pathogens. Nat Immunol. 2008;9(12):1399–406.

17. Cronan MR, Hughes EJ, Brewer WJ, Viswanathan G, Hunt EG, Singh B, et al. A non-canonical type 2 immune response coordinates tuberculous granuloma formation and epithelialization. Cell. 2021;184(7):1757–74.e14.

18. Kramnik I, and Beamer G. Mouse models of human TB pathology: roles in the analysis of necrosis and the development of host-directed therapies. Semin Immunopathol. 2016;38(2):221–37.

19. Florido M, Cooper AM, and Appelberg R. Immunological basis of the development of necrotic lesions following Mycobacterium avium infection. Immunology. 2002;106(4):590–601.

20. Lousada S, Florido M, and Appelberg R. Regulation of granuloma fibrosis by nitric oxide during Mycobacterium avium experimental infection. Int J Exp Pathol. 2006;87(4):307–15.

21. Saunders BM, Dane A, Briscoe H, and Britton WJ. Characterization of immune responses during infection with Mycobacterium avium strains 100, 101 and the recently sequenced 104. Immunology and cell biology. 2002;80(6):544–9.

22. Kondratieva EV, Evstifeev VV, Kondratieva TK, Petrovskaya SN, Pichugin AV, Rubakova EI, et al. I/St Mice Hypersusceptible to Mycobacterium tuberculosis Are Resistant to M. avium. Infection and Immunity. 2007;75(10):4762–8.

23. Verma D, Stapleton M, Gadwa J, Vongtongsalee K, Schenkel AR, Chan ED, et al. Mycobacterium avium Infection in a C3HeB/FeJ Mouse Model. Front Microbiol. 2019;10:693.

24. Kramnik I, Demant P, and Bloom BB. Susceptibility to tuberculosis as a complex genetic trait: analysis using recombinant congenic strains of mice. Novartis Found Symp. 1998;217:120–31; discussion 32-7.

25. Driver ER, Ryan GJ, Hoff DR, Irwin SM, Basaraba RJ, Kramnik I, et al. Evaluation of a mouse model of necrotic granuloma formation using C3HeB/FeJ mice for testing of drugs against Mycobacterium tuberculosis. Antimicrob Agents Chemother. 2012;56(6):3181–95.

26. Harper J, Skerry C, Davis SL, Tasneen R, Weir M, Kramnik I, et al. Mouse model of necrotic tuberculosis granulomas develops hypoxic lesions. The Journal of infectious diseases. 2012;205(4):595–602.

27. Kramnik I, Dietrich WF, Demant P, and Bloom BR. Genetic control of resistance to experimental infection with virulent Mycobacterium tuberculosis. Proceedings of the National Academy of Sciences of the United States of America. 2000;97(15):8560–5.

28. Pan H, Yan BS, Rojas M, Shebzukhov YV, Zhou H, Kobzik L, et al. Ipr1 gene mediates innate immunity to tuberculosis. Nature. 2005;434(7034):767–72.

29. Yan BS, Kirby A, Shebzukhov YV, Daly MJ, and Kramnik I. Genetic architecture of tuberculosis resistance in a mouse model of infection. Genes and immunity. 2006;7(3):201–10.

30. Pichugin AV, Yan BS, Sloutsky A, Kobzik L, and Kramnik I. Dominant role of the sst1 locus in pathogenesis of necrotizing lung granulomas during chronic tuberculosis infection and reactivation in genetically resistant hosts. Am J Pathol. 2009;174(6):2190–201.

31. Sissons J, Yan BS, Pichugin AV, Kirby A, Daly MJ, and Kramnik I. Multigenic control of tuberculosis resistance: analysis of a QTL on mouse chromosome 7 and its synergism with sst1. Genes and immunity. 2009;10(1):37–46.

32. He X, Berland R, Mekasha S, Christensen TG, Alroy J, Kramnik I, et al. The sst1 resistance locus regulates evasion of type I interferon signaling by Chlamydia pneumoniae as a disease tolerance mechanism. PLoS Pathog. 2013;9(8):e1003569.

33. Bhattacharya B, Xiao S, Chatterjee S, Urbanowski ME, Ordonez AA, Ihms EA, et al. The integrated stress response mediates necrosis in murine Mycobacterium tuberculosis granulomas. J Clin Invest. 2020.

34. Brownhill E, Yabaji SM, Zhernovkov V, Rukhlenko OS, Seidel K, Bhattacharya B, et al. Maladaptive oxidative stress cascade drives type I interferon hyperactivity in TNF activated macrophages promoting necrosis in murine tuberculosis granulomas. bioRxiv. 2020:2020.12.14.422743.

35. Ji DX, Yamashiro LH, Chen KJ, Mukaida N, Kramnik I, Darwin KH, et al. Type I interferon-driven susceptibility to Mycobacterium tuberculosis is mediated by IL-1Ra. Nat Microbiol. 2019;4(12):2128–35.

36. Boyartchuk V, Rojas M, Yan BS, Jobe O, Hurt N, Dorfman DM, et al. The host resistance locus sst1 controls innate immunity to Listeria monocytogenes infection in immunodeficient mice. Journal of immunology. 2004;173(8):5112–20.

37. Ji DX, Witt KC, Kotov DI, Margolis SR, Louie A, Chevée V, et al. Role of the transcriptional regulator SP140 in resistance to bacterial infections via repression of type I interferons. bioRxiv. 2021.

38. Larsen SE, Reese VA, Pecor T, Berube BJ, Cooper SK, Brewer G, et al. Subunit vaccine protects against a clinical isolate of Mycobacterium avium in wild type and immunocompromised mouse models. Scientific Reports. 2021;11(1):9040.

39. Scheu S, Dresing P, and Locksley RM. Visualization of IFNβ production by plasmacytoid versus conventional dendritic cells under specific stimulation conditions in vivo. Proceedings of the National Academy of Sciences of the United States of America. 2008;105(51):20416–21.

40. Scriba TJ, Penn-Nicholson A, Shankar S, Hraha T, Thompson EG, Sterling D, et al. Sequential inflammatory processes define human progression from M. tuberculosis infection to tuberculosis disease. PLoS pathogens. 2017;13(11):e1006687.

41. McNab F, Mayer-Barber K, Sher A, Wack A, and O&Garra A. Type I interferons in infectious disease. Nature Reviews Immunology. 2015;15(2):87–103.

42. Moreira-Teixeira L, Mayer-Barber K, Sher A, and O&Garra A. Type I interferons in tuberculosis: Foe and occasionally friend. The Journal of experimental medicine. 2018;215(5):1273–85.

43. Andréjak C, Almeida DV, Tyagi S, Converse PJ, Ammerman NC, and Grosset JH. Characterization of mouse models of Mycobacterium avium complex infection and evaluation of drug combinations. Antimicrobial agents and chemotherapy. 2015;59(4):2129–35.

44. Appelberg R. Pathogenesis of Mycobacterium avium infection: typical responses to an atypical mycobacterium? Immunol Res. 2006;35(3):179–90.

45. Hwang JA, Kim S, Jo K-W, and Shim TS. Natural history of Mycobacterium avium complex lung disease in untreated patients with stable course. The European respiratory journal. 2017;49(3).

46. Yan BS, Pichugin AV, Jobe O, Helming L, Eruslanov EB, Gutierrez-Pabello JA, et al. Progression of pulmonary tuberculosis and efficiency of bacillus Calmette-Guerin vaccination are genetically controlled via a common sst1-mediated mechanism of innate immunity. Journal of immunology. 2007;179(10):6919–32.

47. Jabado N, Jankowski A, Dougaparsad S, Picard V, Grinstein S, and Gros P. Natural resistance to intracellular infections: natural resistance-associated macrophage protein 1 (Nramp1) functions as a pH-dependent manganese transporter at the phagosomal membrane. J Exp Med. 2000;192(9):1237–48.

48. GOMES, and APPELBERG. Evidence for a link between iron metabolism and Nramp1 gene function in innate resistance against Mycobacterium avium. Immunology. 1998;95(2):165–8.

49. Linge I, Petrova E, Dyatlov A, Kondratieva T, Logunova N, Majorov K, et al. Reciprocal control of Mycobacterium avium and Mycobacterium tuberculosis infections by the alleles of the classic Class II H2-Aβ gene in mice. Infection, genetics and evolution: journal of molecular epidemiology and evolutionary genetics in infectious diseases. 2019;74:103933.

50. Pessanha AP, Martins RA, Mattos-Guaraldi AL, Vianna A, and Moreira LO. Arginase-1 expression in granulomas of tuberculosis patients. FEMS Immunol Med Microbiol. 2012;66(2):265–8.

51. Heitmann L, Abad Dar M, Schreiber T, Erdmann H, Behrends J, Mckenzie AN, et al. The IL-13/IL-4R αaxis is involved in tuberculosis-associated pathology. The Journal of pathology. 2014;234(3):338–50.

52. MacMicking JD, North RJ, LaCourse R, Mudgett JS, Shah SK, and Nathan CF. Identification of nitric oxide synthase as a protective locus against tuberculosis. Proceedings of the National Academy of Sciences of the United States of America. 1997;94(10):5243–8.

53. Duque-Correa MA, Kuhl AA, Rodriguez PC, Zedler U, Schommer-Leitner S, Rao M, et al. Macrophage arginase-1 controls bacterial growth and pathology in hypoxic tuberculosis granulomas. Proceedings of the National Academy of Sciences of the United States of America. 2014;111(38):E4024–32.

54. Gomes MS, Florido M, Pais TF, and Appelberg R. Improved clearance of Mycobacterium avium upon disruption of the inducible nitric oxide synthase gene. Journal of immunology. 1999;162(11):6734–9.

55. Bronte V, Serafini P, De Santo C, Marigo I, Tosello V, Mazzoni A, et al. IL-4-induced arginase 1 suppresses alloreactive T cells in tumor-bearing mice. The Journal of Immunology. 2003;170(1):270–8.

56. Murray PJ, Allen JE, Biswas SK, Fisher EA, Gilroy DW, Goerdt S, et al. Macrophage activation and polarization: nomenclature and experimental guidelines. Immunity. 2014;41(1):14–20.

57. Kramnik I, Radzioch D, and Skamene E. T-helper 1-like subset selection in Mycobacterium bovis bacillus Calmette-Guérin-infected resistant and susceptible mice. Immunology. 1994;81(4):618–25.

58. Darrah PA, Zeppa JJ, Maiello P, Hackney JA, Wadsworth II MH, Hughes TK, et al. Prevention of tuberculosis in macaques after intravenous BCG immunization. Nature. 2020;577(7788):95–102.

59. Greenberg A, Huber BR, Liu DX, Logue JP, Hischak AMW, Hart RJ, et al. Quantification of Viral and Host Biomarkers in the Liver of Rhesus Macaques: A Longitudinal Study of Zaire Ebolavirus Strain Kikwit (EBOV/Kik). Am J Pathol. 2020;190(7):1449–60.

60. Shanmugasundaram U, Bucsan AN, Ganatra SR, Ibegbu C, Quezada M, Blair RV, et al. Pulmonary Mycobacterium tuberculosis control associates with CXCR3- and CCR6-expressing antigen-specific Th1 and Th17 cell recruitment. JCI Insight. 2020;5(14).

61. Gideon HP, Hughes TK, Wadsworth II MH, Tu AA, Gierahn TM, Hopkins FH, et al. Single-cell profiling of tuberculosis lung granulomas reveals functional lymphocyte signatures of bacterial control. 2020:1–60.

62. Blanc L, Lenaerts A, Dartois V, and Prideaux B. Visualization of Mycobacterial Biomarkers and Tuberculosis Drugs in Infected Tissue by MALDI-MS Imaging. Anal Chem. 2018;90(10):6275–82.

63. Prideaux B, Lenaerts A, and Dartois V. Imaging and spatially resolved quantification of drug distribution in tissues by mass spectrometry. Curr Opin Chem Biol. 2018;44:93–100.

