## Supplemental Table 1 for "Dissemination and progression of pulmonary *Mycobacterium avium* infection in mouse model is associated with type 2 macrophage activation"

| Supplementary Table 1. Pulmonary Histopathology Ordinal Score Summary (intranasal inoculation) | | | | | |
|---|---|---|---|---|---|
| Weeks post-infection and inoculation dosage | Animal ID | Slide # | Ordinal Score | strain | sex |
| 4 wks 10^6 | C1-1 | 1 | 0 | B6.Sst1S | F |
| | | 4 | 1 | | |
| | | 7 | 1 | | |
| | C1-2 | 1 | 1 | B6.Sst1S | F |
| | | 4 | 1 | | |
| | | 7 | 1 | | |
| | C1-3 | 1 | 0 | B6.Sst1S | M |
| | | 4 | 1 | | |
| | | 7 | 1 | | |
| | C5-1 | 1 | 0 | B6.Sst1S | M |
| | | 4 | 0 | | |
| | | 7 | 0 | | |
| | C5-2 | 1 | 1 | B6.Sst1S | M |
| | | 4 | 1 | | |
| | | 7 | 1 | | |
| | C5-3 | 1 | 1 | B6.Sst1S | M |
| | | 4 | 2 | | |
| | | 7 | 2 | | |
| 6 wks 10^7 | 12-1 | 1 | 1 | B6.Sst1S | F |
| | | 4 | 2 | | |
| | | 7 | 2 | | |
| | 12-2 | 1 | 2 | B6.Sst1S | F |
| | | 4 | 2 | | |
| | | 7 | 2 | | |
| | 13-1 | 1 | 1 | B6.Sst1S | F |
| | | 4 | 1 | | |
| | | 7 | 1 | | |
| | 13-2 | 1 | 1 | B6.Sst1S | F |
| | | 4 | 2 | | |
| | | 7 | 1 | | |
| 8 wks 10^6 | 4-1 | 1 | 2 | B6.Sst1S | M |
| | | 4 | 2 | | |
| | | 7 | 2 | | |

| | | | | | |
|---|---|---|---|---|---|
| | 4-2 | 1 | 1 | B6.Sst1S | M |
| | | 4 | 2 | | |
| | | 7 | 2 | | |
| 10 wks 10^6 | 2-1 | 1 | 2 | B6.Sst1S | F |
| | | 4 | 2 | | |
| | | 7 | 2 | | |
| | 2-2 | 1 | 1 | B6.Sst1S | F |
| | | 4 | 2 | | |
| | | 7 | 2 | | |
| | 3-1 | 1 | 2 | B6.Sst1S | F |
| | | 4 | 2 | | |
| | | 7 | 2 | | |
| 10 wks 10^7 | 13-3 | 1 | 1 | B6.Sst1S | F |
| | | 4 | 1 | | |
| | | 7 | 1 | | |
| | 14-1 | 1 | 2 | B6.Sst1S | F |
| | | 4 | 2 | | |
| | | 7 | 2 | | |
| 12 wks 10^6 | 3-2 | 1 | 2 | B6.Sst1S | F |
| | | 4 | 2 | | |
| | | 7 | 2 | | |
| | 3-3 | 1 | 2 | B6.Sst1S | F |
| | | 4 | 2 | | |
| | | 7 | 2 | | |
| | 3-4 | 1 | 2 | B6.Sst1S | F |
| | | 4 | 2 | | |
| | | 7 | 2 | | |
| | 14-2 | 1 | 2 | B6.Sst1S | F |
| | | 4 | 3 | | |
| | | 7 | 2 | | |
| | 14-3 | 1 | 2 | B6.Sst1S | F |
| | | 4 | 2 | | |
| | | 7 | 2 | | |
