## Supplemental Table 2 for "Dissemination and progression of pulmonary *Mycobacterium avium* infection in mouse model is associated with type 2 macrophage activation"

| Supplementary Table 2. Pulmonary Histopathology Ordinal Score Summary (Left mainstream intrabronchial inoculation) | | | | | |
|---|---|---|---|---|---|
| Weeks Post-Infection (WPI) | Animal ID | Depth of Block (mm from ventral surface) | Strain | Left lung (inoculation site) | Right lungs (un-inoculated) |
| 12 WPI | 1 | 1 | WT | 2 | 2 |
| | | 2 | | 2 | 2 |
| | 2 | 1 | | 1 | 1 |
| | | 2 | | 1 | 1 |
| | 3 | 1 | | 1 | 1 |
| | | 2 | | 1 | 1 |
| | 4 | 1 | | 2 | 1 |
| | | 2 | | 2 | 1 |
| | 13 | 1 | B6.Sst1S | 2 | 2 |
| | | 2 | | 2 | 2 |
| | 14 | 1 | | 1 | 0 |
| | | 2 | | 1 | 1 |
| | 15 | 1 | | 0 | 0 |
| | | 2 | | 0 | 0 |
| 16 WPI | 5 | 1 | WT | 3 | 2 |
| | | 2 | | 3 | 2 |
| | 6 | 1 | | 3 | 2 |
| | | 2 | | 3 | 2 |
| | 7 | 1 | | 2 | 2 |
| | | 2 | | 2 | 1 |
| | 8 | 1 | | 2 | 2 |
| | | 2 | | 2 | 2 |
| | 16 | 1 | B6.Sst1S | 3 | 3 |
| | | 2 | | 3 | 3 |
| | 17 | 1 | | 3 | 2 |
| | | 2 | | 3 | 2 |
| | 18 | 1 | | 0 | 0 |
| | | 2 | | 0 | 0 |

| | | | | | |
|---|---|---|---|---|---|
| | 19 | 1 | | 2 | 2 |
| | | 2 | | 2 | 2 |
| 21 wks 10^5 M avium IT | 9 | 1 | WT | 2 | 2 |
| | | 2 | | 2 | 2 |
| | | 3 | | 2 | 2 |
| | 10 | 1 | | 3 | 3 |
| | | 2 | | 3 | 3 |
| | | 3 | | 3 | 3 |
| | 11 | 1 | | 2 | 3 |
| | | 2 | | 3 | 3 |
| | | 3 | | 3 | 3 |
| | 12 | 1 | | 0 | 0 |
| | | 2 | | 0 | 0 |
| | | 3 | | 0 | 0 |
| | 20 | 1 | B6.Sst1S | 3 | 3 |
| | | 2 | | 3 | 3 |
| | | 3 | | 3 | 2 |
| | 21 | 1 | | 3 | 2 |
| | | 2 | | 3 | 2 |
| | | 3 | | 3 | 2 |
| | 22 | 1 | | 3 | 2 |
| | | 2 | | 3 | 3 |
