## Supplementary Figure Legends for "Dissemination and progression of pulmonary *Mycobacterium avium* infection in mouse model is associated with type 2 macrophage activation"

**Supplemental figure legends**

**Supplemental Figure 1.** Representative large (left panel) and small (right panel) secondary lesions as identified by a random forest classifier that detected iNOS, Arg1, M. *avium* (M. *av*), and interspersed CD3+ cells in the uninoculated lung lobes of B6.Sst1S mice at 16 wpi intrabronchial infected with Mycobacterium *avium* (M. *av).* Large secondary lesions represented the top 10 percentile lesions by area for each given specimen. Small secondary lesions represented lesions between 50-60th percentile lesions by area for each given specimen. Lesions were excluded if they were directly affiliated with a bronchiole.

**Supplemental Figure 2.** Comparison between primary lesions and secondary lesions following unilateral left mainstream intrabronchial inoculation at 12- (A-F), 16- (G-L) and 21- (M-R) weeks post-infection (wpi). **A:** Histopathologic scores. **B:** Lesion area. **C:** M. *av* immunoreactivity area quantification (AQ). **D:** iNOS AQ. **E:** Arg1 AQ. **F:** T cell density. **G:** Histopathologic scores. **H:** Lesion area. **I:** M. *av* AQ. **J:** iNOS AQ. **K:** Arg1 AQ. **L:** T cell density. **M:** Histopathologic scores. **N:** Lesion area. **O:** M. *av* AQ. **P:** iNOS AQ. **Q:** Arg1 AQ.

**R:** CD3+ AQ. For A, G, and M, each data point represents a single tissue section. For B, H, and N, each data point represents a single lesion. For C-F, I-L, O-R, each data point represents a single animal. Data are expressed as means ± SD or with Tukey whiskers. Asterisks

denote P values with Mann Whitney test (*) or Student’s unpaired T test (*-*).

∗P < 0.05, ∗∗P < 0.005, ∗∗∗P < 0.0005, and ∗∗∗∗P

**Supplemental Figure 3** B6 WT vs. B6.Sst1S ordinal histopathology comparisons following unilateral left mainstream intrabronchial inoculation at 12, 16- and 21-weeks post-infection (wpi) and the impacts of *sst1*-S in small secondary lesions at 16 wpi. **A:** Histopathology scores at 12 wpi. **B:** Histopathology scores at 16 wpi. **C:** Histopathology scores at 21 wpi. **D:** Representative raw fluorescent multiplex immunohistochemical (mIHC) image of B6 WT small secondary lesion. **E:** Representative raw fluorescent mIHC image of B6.Sst1S small secondary lesion. **F-N**: Comparison of fluorescent mIHC results in B6 WT and B6.Sst1S small secondary lesions; **F.** iNOS AQ, **G** iNOS+ Arg1- M. avium- AQ, **H.** iNOS+ Arg1- M. avium+ AQ, **I.** Arg1 AQ, **J.** Arg1+ iNOS- M. avium- AQ, **K.** Arg1+ iNOS- M. avium+ AQ, **L.** Arg1+ iNOS+ M. avium+ AQ, **M.** Arg1+ iNOS+ M. avium- AQ, **N.** T cell lesion density in B6 WT and B6.Sst1S small secondary lesions. For A-C, each data point represents one tissue section. For F-M, percentage area immunoreactivity was calculated in relation to area of given lesion. Each data point represents one lesion. Data are expressed as means ± SD or with Tukey whiskers. Asterisks denote P values with Mann Whitney test (*) or Student’s unpaired T test (*-*). ∗P < 0.05, ∗∗P < 0.005, ∗∗∗P < 0.0005, and ∗∗∗∗P. Original magnification, 100x (**D** and **E**).
