## Supplementary Figures for "Dissemination and progression of pulmonary *Mycobacterium avium* infection in mouse model is associated with type 2 macrophage activation"

### Slide 1
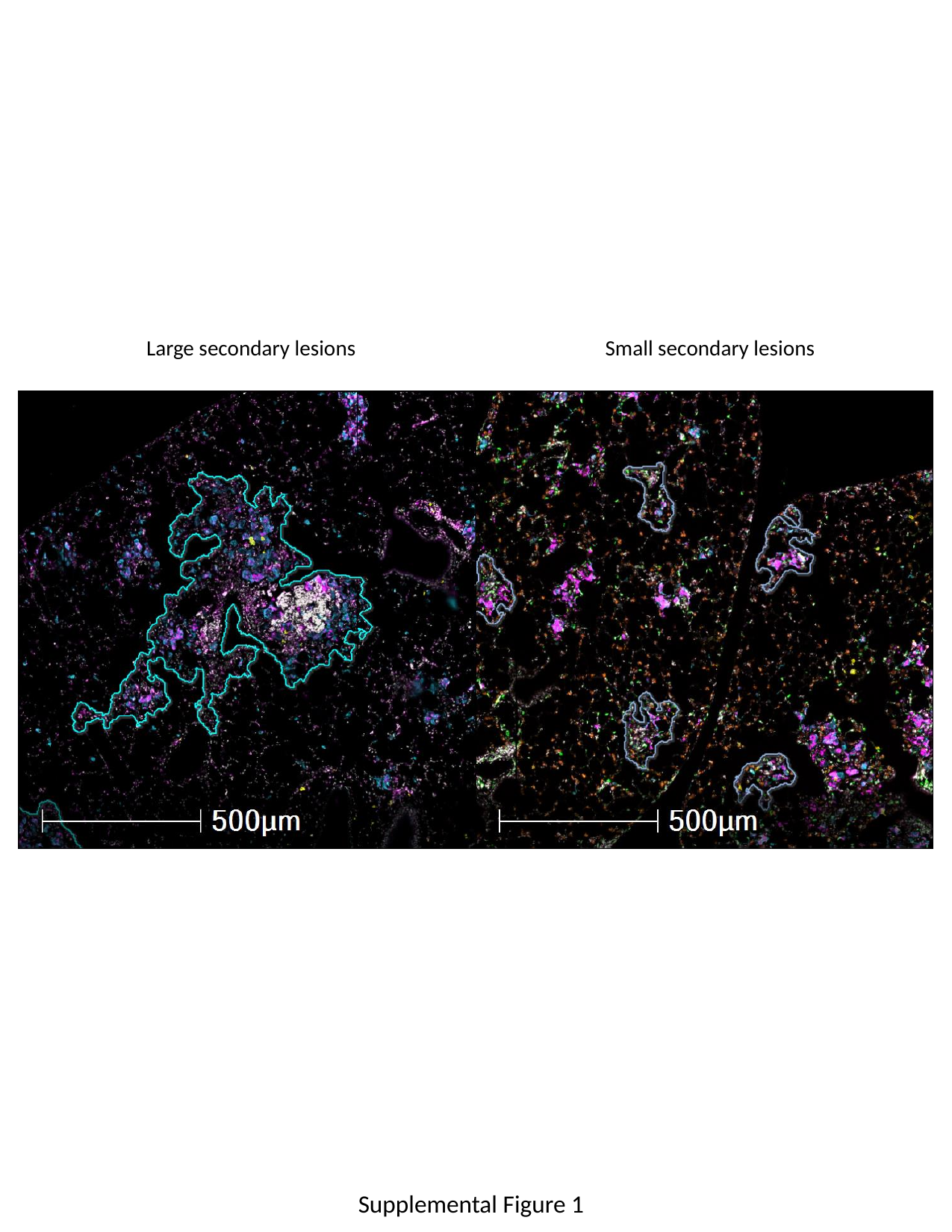

Large secondary lesions
Small secondary lesions
Supplemental Figure 1

### Slide 2
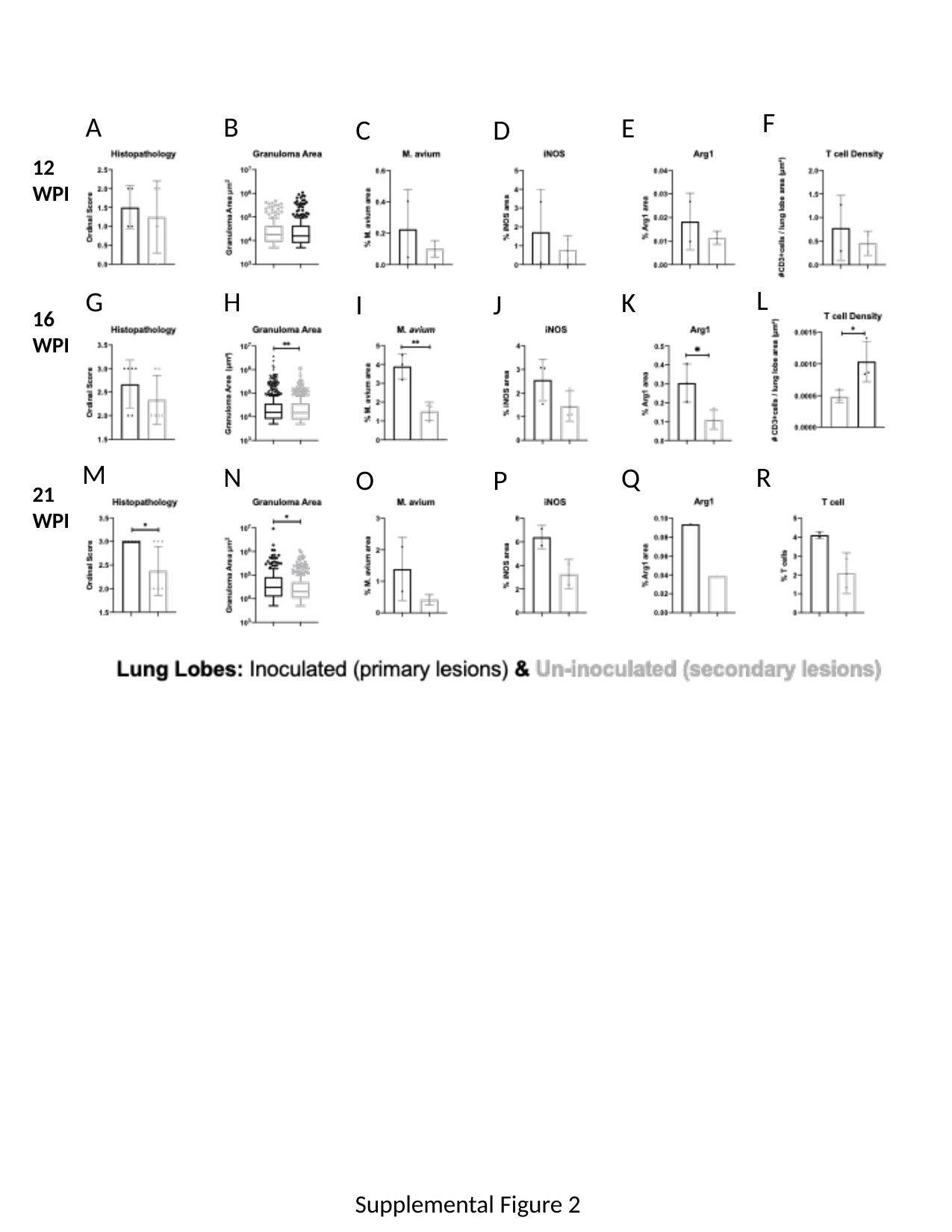

F
A
B
E
C
D
12 WPI
L
G
H
K
I
J
16 WPI
M
N
R
Q
O
P
21 WPI
Supplemental Figure 2

### Slide 3
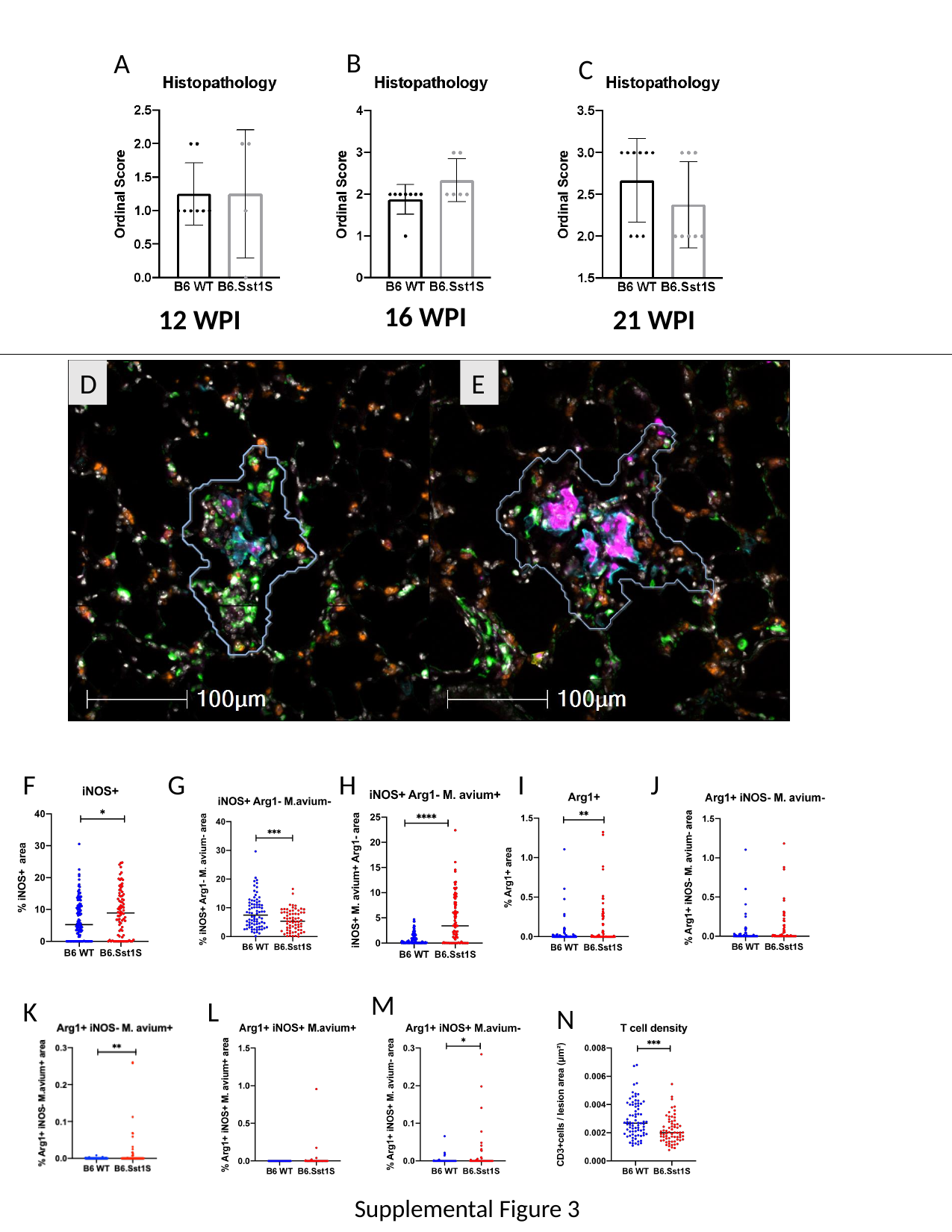

B
A
C
16 WPI
12 WPI
21 WPI
D
E
I
J
G
F
H
M
L
K
N
Supplemental Figure 3
